# Neighbourhood species richness and drought-tolerance traits modulate tree growth and δ^13^C responses to drought

**DOI:** 10.1101/2022.11.22.517351

**Authors:** Florian Schnabel, Kathryn E. Barry, Susanne Eckhardt, Joannès Guillemot, Heike Geilmann, Anja Kahl, Heiko Moossen, Jürgen Bauhus, Christian Wirth

## Abstract

Mixed-species forests are promoted as a forest management strategy for climate change adaptation, but whether they are more resistant to drought than monospecific forests remains contested. Particularly, the trait-based mechanisms driving the role of tree diversity under drought remain elusive.
Using tree cores from a large-scale biodiversity experiment, we investigated tree growth and physiological stress responses (i.e. increase in wood carbon isotopic ratio; δ^13^C) to changes in climate-induced water availability (wet to dry years) along gradients in neighbourhood tree species richness and drought-tolerance traits. We hypothesized that neighbourhood species richness increases growth and decreases δ^13^C and that these relationships are modulated by the abiotic (i.e. climatic conditions) and the biotic context. We characterized the biotic context using drought-tolerance traits of focal trees and their neighbours. These traits are related to cavitation resistance vs resource acquisition and stomatal control.
Tree growth increased with neighbourhood species richness. However, we did not observe a universal relief of water stress in species-rich neighbourhoods. Neighbourhood species richness effects on growth and δ^13^C did not strengthen from wet to dry years. Instead, richness-growth and richness-δ^13^C relationships were modulated by climatic conditions and the traits of trees and their neighbours. At either end of each drought-tolerance gradient, species responded in opposing directions during drought and non-drought years.
We show that species’ drought-tolerance traits can explain the strength and nature of biodiversity-ecosystem functioning relationships in experimental tree communities experiencing drought. Mixing tree species can increase growth but may not universally relieve drought stress.

**One-sentence summary:** The drought-tolerance traits of trees and their neighbours determine biodiversity-ecosystem functioning relationships in experimental tree communities.

## Introduction

The world’s forests are experiencing widespread mortality events due to climate extremes like droughts (Hartmann *et al*., 2022). Such climate extremes are predicted to increase in frequency and intensity with climate change in many regions (IPCC, 2022), threatening multiple ecosystem services that forests provide, including their capacity to mitigate climate change through carbon sequestration and storage (Anderegg *et al*., 2020). Large-scale forest restoration initiatives such as the Bonn Challenge, which aims to restore 350 Mha of forests by 2030 to mitigate climate change (Brancalion *et al*., 2019), need to optimize productivity and thus carbon storage while at the same time increasing these restored forests’ ecological stability against climate extremes (Bauhus *et al*., 2017). One key management strategy suggested to achieve this desired synergy between productivity and stability is to establish and maintain tree species mixtures instead of monocultures (Jucker *et al*., 2014; Messier *et al*., 2021).

There is now accumulating evidence that species-rich forests provide higher levels and higher stability of various ecosystem functions than species-poor or monospecific forests (Messier *et al*., 2021; Schnabel *et al*., 2021; van der Plas, 2019). Such tree species richness effects can be best studied at the neighbourhood scale, where tree-tree interactions occur (Trogisch *et al*., 2021). This is because neighbourhood analyses allow studying neighbourhood species richness (hereafter called ‘NSR’) effects alongside factors such as tree size and competition, which can impact tree performance (Forrester & Pretzsch, 2015; Stoll & Newbery, 2005). Importantly, positive NSR effects, for instance on tree growth, may be more pronounced under stress (Bertness & Callaway, 1994; Forrester & Bauhus, 2016) and have been shown to increase in dry compared to wet years (Fichtner *et al*., 2020; Schnabel *et al*., 2019). However, studies on net tree mixture responses to drought have produced divergent results, including positive, neutral and negative diversity effects under drought (reviewed by Grossiord (2020)) and we thus do not yet know when diversity is beneficial for forest functioning under drought. These mixed results may be driven by species-specific water-use strategies. This is because water-use strategies, which can be studied using functional traits, impact how species respond to their surrounding abiotic (i.e. climatic conditions) and biotic environment (i.e. their tree neighbours) (Forrester, 2017). We characterized this biotic context, which has received little attention compared to the many studies that examined the abiotic context dependency of biodiversity-ecosystem functioning (BEF) relationships (e.g. Forrester & Bauhus, 2016; Grossiord, 2020; Jucker *et al*., 2016), as traits of focal trees and as traits of their neighbours.

The recent history of an abiotic and biotic control on BEF relationships can be analysed in the tree ring record of a focal tree, which captures its response to neighbours and climatic variation (Schweingruber, 1996; Vitali *et al*., 2018). The width of annual growth rings is an indicator of a tree’s reaction to climate, with reduced growth in dry compared to wet years indicating increased drought stress (Schwarz *et al*., 2020). In addition to growth, the isotopic ratio of ^13^C/^12^C in wood (hereafter ‘δ^13^C’) is a principal indicator of a tree’s physiological reaction to water limitation and drought, with higher δ^13^C values suggesting greater drought stress. δ^13^C captures the ratio of CO_2_ concentration inside the cells and in the atmosphere during the time of wood formation, thus reflecting the balance between net CO_2_ assimilation and stomatal conductance of trees (Farquhar *et al*., 1989; Grams *et al*., 2007). Under drought stress, trees close their stomata to avoid water loss from transpiration, and as stomatal conductance is more affected than assimilation, δ^13^C increases under drought (Farquhar *et al*., 1989; Grossiord *et al*., 2014). Therefore, δ^13^C is widely recognized as an indicator of the severity of drought exposure in trees, and we consider it as a proxy for neighbourhood-scale water availability (Grossiord *et al*., 2014; Jucker *et al*., 2017).

Among other relevant traits, two key traits proposed to influence tree responses to drought are cavitation resistance and the stringency of stomatal control (McDowell *et al*., 2008), which we collectively refer to as ‘drought-tolerance traits’ (see Schnabel *et al*., 2021 for details). In brief, xylem resistance to cavitation reduces embolism risk in vessels, which impair water transport and, at advanced stages, induce desiccation and, ultimately, tree death (Choat *et al*., 2012). Cavitation resistance is often quantified as the water potential where 50% of conductivity is lost due to cavitation (Ψ_50_; Choat *et al*., 2012). Moreover, cavitation resistance has been shown to be associated with classic traits of the leaf economics spectrum in tropical tree species (Guillemot *et al*., 2022) and thus with resource use strategies. Accordingly, cavitation-sensitive species often have traits indicative of acquisitive resource use (Fichtner *et al*., 2020; Schnabel *et al*., 2021). In addition to cavitation resistance, stomatal control differs among species: some species close their stomata early during water shortages to avoid transpirational water loss (called ‘water-saving’ or ‘isohydric’ species), whereas others keep their stomata open despite increasingly negative water potentials and increasing cavitation risks (called ‘water-spending’ or ‘anisohydric’ species) (Martínez-Vilalta & Garcia-Forner, 2017; McDowell *et al*., 2008). In line with current perspectives, we do not use the regulation of leaf water potential (Ψ_L_) as measure for stomatal control, as it was shown to be a poor proxy (Martínez-Vilalta & Garcia-Forner, 2017). Instead, we use physiological traits such as stomatal conductance and its control under increasing vapour pressure deficits (VPD) to quantify stomatal control as a gradient from water savers, which close their stomata early as water stress develops, to water spenders, which keep their stomata open despite increasing VPD (Kröber *et al*., 2014; Schnabel *et al*., 2021). Diversity in these traits, hereafter referred to as ‘resistance-acquisition’ and ‘stomatal control’ traits, has been recently shown to be positively related to the stability of forest community productivity under highly variable climatic conditions (Schnabel *et al*., 2021). However, these traits have not been used in a comprehensive framework to characterize the functional identity of focal trees (i.e. their traits) and their neighbours (i.e. the neighbourhood mean values of traits) to understand how these functional identities influence growth and wood δ^13^C responses to the interactive effects of NSR and contrasting climatic conditions (such as particularly dry and wet years).

A focal trees’ functional (trait) identity, hereafter ‘focal tree traits’, may be crucial to understand responses of tree growth and δ^13^C to the interactive effects of drought and NSR. Consistent with this expectation, Fichtner *et al*. (2020) showed that positive NSR effects on aboveground wood production were strongest for cavitation-sensitive species during drought. Moreover, another study, even though conducted in a wet year, found that increasing acquisitiveness and NSR caused decreased δ^13^C values in tree twig tissues, indicating enhanced water availability in diverse neighbourhoods (Jansen *et al*., 2021). In this context, growth may be more strongly related to resistance-acquisition traits and thus the leaf economics spectrum (Reich, 2014). Alternatively, wood δ^13^C may be primarily controlled by stomata aperture (Farquhar *et al*., 1989) and thus stomatal control traits.

The trait identity of a focal trees’ neighbours, hereafter ‘neighbour traits’, may also influence the growth and δ^13^C of focal trees through their influence on biotic interactions (e.g. Fortunel *et al*., 2016; Trogisch *et al*., 2021; Yang *et al*., 2021). In this view, neighbour traits may alter water use and with this local water availability. For example, during drought, growth reductions in water spenders may be lower and δ^13^C increases smaller when growing with more water-saving neighbours because the reduced stomatal conductance of the latter may decrease overall water consumption and thus drought stress in focal trees (Forrester, 2017). Conversely, being surrounded by water spending neighbours during drought may amplify focal tree growth reductions and δ^13^C increases.

Here, we aim to understand how drought-tolerance traits influence the relationship between NSR and growth, and NSR and δ^13^C under variable climatic conditions to shed light on the abiotic and biotic context dependency of forest BEF relationships. We use trait-based neighbourhood models that account for NSR as well as focal tree and neighbour traits and examine how they jointly influence focal tree growth and δ^13^C in a climatically dry, normal and wet year in a large-scale sub-tropical tree biodiversity-ecosystem functioning experiment (BEF-China experiment; Bruelheide *et al*., 2014; Huang *et al*., 2018). Specifically, we tested the following hypothesis:

**H1**: NSR increases growth and decreases δ^13^C of focal trees, and the strength of this diversity effect increases from wet to dry years.
**H2**: Drought-tolerance traits of focal trees determine the relationship between NSR and growth, and NSR and δ^13^C under variable climatic conditions. Specifically, during drought, NSR increases growth and decreases δ^13^C for acquisitive and water-spending species, while the reverse pattern is found for cavitation-resistant and water-saving species.
**H3**: Drought-tolerance traits of neighbours influence the effect of climate on focal tree growth. Specifically, during drought, acquisitive and water-spending neighbours amplify drought stress.

## Materials and Methods

### Study site and experimental design

We sampled trees in a large-scale tree biodiversity-ecosystem functioning experiment located in Xingangshan, Dexing, Jiangxi province, China (29°08′N to 29°11′N, 117°90′E to 117°93′E), the BEF-China experiment (Bruelheide *et al*., 2014; Huang *et al*., 2018). The experiment has two sites: A and B, each approximately 20 ha in size. The sites are characterised by a subtropical, monsoon climate with a mean annual temperature of 16.7 °C and a yearly precipitation sum of 1821 mm (mean from 1971–2000; Yang *et al*., 2013), with distinct differences between seasons. Summers are humid, with most annual precipitation falling from April to July, while winters are drier and cold (Gheyret *et al*., 2021). Deciduous and evergreen broadleaved trees dominate the diverse native forests in the study region, sometimes mixed with conifers (Bruelheide *et al*., 2014). The regions location between tropical and temperate climates with their respective flora (Shi *et al*., 2014; Wang *et al*., 2007), makes it ideal for studying diverse water-use strategies and species responses to variable climatic conditions (Schnabel *et al*., 2021). Based on a pool of 40 native evergreen and deciduous broadleaf tree species, experimental tree species richness gradients were created with monocultures 2-, 4-, 8-, 16- and 24-species mixtures. Species were assigned to different extinction scenarios following a broken-stick design, ensuring that all species were represented at each species richness level (Bruelheide *et al*., 2014; Huang *et al*., 2018). In 2009 (site A) and 2010 (site B), overall, 226,400 individual trees were planted at a distance of 1.29 meters on plots with a size of 25.8 × 25.8 m^2^, with 400 trees being planted per plot. Species compositions and tree positions within plots were randomly assigned to each plot.

### Climate-based selection of study years

We selected three study years with contrasting climatic conditions, a comparably wet (2016), an intermediate (2017) and a particularly dry year (2018). The analysis of consecutive years allowed us to minimise other factors than climate that may influence growth and δ^13^C, such as changes in stand structure. We used the standardised precipitation evapotranspiration index (SPEI) (Vicente-Serrano *et al*., 2010) calculated from a high-resolution time-series of interpolated climate station data (CRU TS v4.04; Harris *et al*., 2020) to characterise climatic conditions. The SPEI represents a standardised climatic water balance of precipitation minus potential evapotranspiration. We selected study years following suggestions by Schwarz *et al*. (2020) by comparing SPEI series for the peak vegetation period (SPEI3, April-July), the full vegetation period (SPEI6, April-September) and the entire year (SPEI12, October-September), using 1901-2019 as climate reference period (Fig. 1A; Fig. S1-S2). The subtropical vegetation period in the study region ranges from April-September with peak growth at the end of April (Gheyret *et al*., 2021), which corresponds well with the selected SPEI lengths. All periods (SPEI3, SPEI6, SPEI12) showed the same pattern of decreasing SPEI values from 2016-2017- 2018 (Fig. S1), with drought severity in the dry year being comparable to drought conditions in the last 40 years (Fig. S2). In addition, we also examined intra-annual and non-standardized climatic water balances (Fig. S3).

**Fig. 1.**
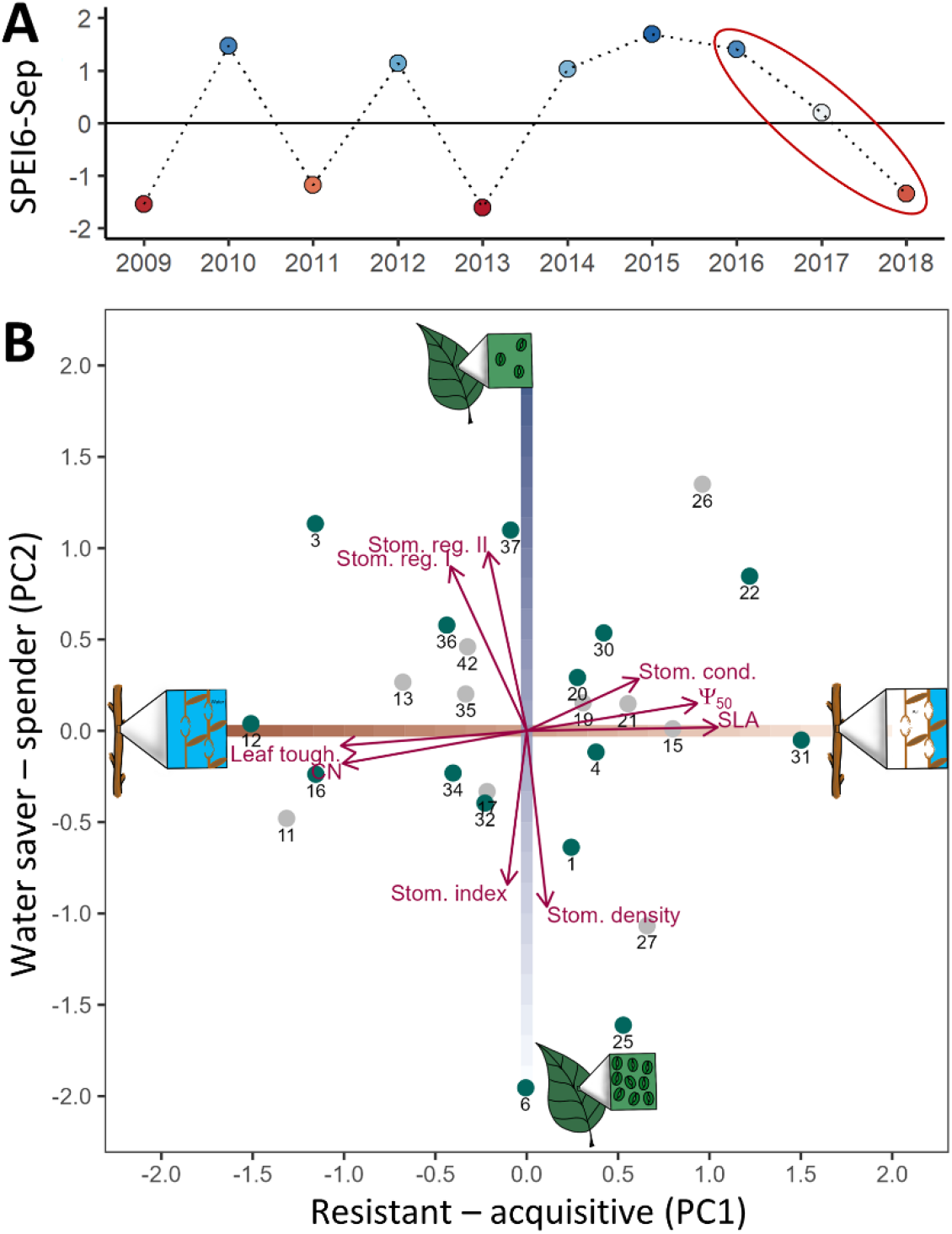
Selection of study years and species. **(A)** Climate-based selection of the study years 2016-2018 based on the standardised precipitation evapotranspiration index of the vegetation period (SPEI6, April-September). Blue points indicate wetter and red points drier conditions compared to the long-term mean; values below -1 and above 1 can be considered exceptional. **(B)** Species selection via their drought-tolerance traits based on principal component analysis (PCA) adapted from Schnabel *et al*. (2021); species represented by codes. PC1 reflects a resistance-acquisition gradient running from cavitation-resistant (low Ψ_50_, high leaf toughness and C/N ratio) to acquisitive species (high Ψ_50_, specific leaf area (SLA), maximum stomatal conductance). PC2 reflects a stomatal control gradient running from water spenders with late stomata closure under decreasing vapour pressure deficits (stomatal regulation traits I and II) to water savers with fast stomata closure (high stomatal density and stomatal index). The sketches illustrate the trait gradients: (PC1) high *vs* low cavitation resistance, (PC2) few vs abundant stomata. We selected focal trees from 15 species to cover the trait space (highlighted in green). See Table S1 for a list of tree species and Table S2 for details on the traits.

### Species selection via drought-tolerance traits

We used this experimental set-up to select tree species along two orthogonal trait gradients related to resistance-acquisition and stomatal control traits (Fig. 1B) which allowed us to study their relative contributions to tree growth and δ^13^C. For this purpose, we relied on species-specific trait data related to cavitation resistance, resource acquisitiveness and stomatal control measured in the experiment (Table S2; Kröber *et al*., 2014; Kröber & Bruelheide, 2014). Trait data were analysed with principal component analysis (PCA), which partitioned the variation in drought-tolerance traits into two orthogonal trait gradients, a resistance-acquisition (PC1) and stomatal control (PC2) trait gradient (see Schnabel *et al*. (2021) for details). In brief, we quantified cavitation resistance as the water potential (Ψ_50_) at which 50% of xylem conductivity is lost due to cavitation, which is a key physiological trait to characterise a species drought tolerance (Choat *et al*., 2012). In our study system, Ψ_50_ is related to classic traits of the leaf economics spectrum (Reich, 2014) in that cavitation-resistant species (low Ψ_50_ values) are also characterised by traits indicative of conservative resource use (tough leaves and high C/N ratio), hereafter referred to as ‘resistant species’ (Fig. 1B). In contrast, cavitation-sensitive species (less negative Ψ_50_ values) have traits indicative of acquisitive resource use such as high specific leaf area (SLA) and high maximum stomatal conductance (g_smax_), hereafter referred to as ‘acquisitive species’. Including this gradient provided a balanced selection of deciduous and evergreen species. Second, we quantified stomatal control using modelled curves of stomatal conductance (g_s_) under increasing vapour pressure deficits (VPD) and morphological traits (stomatal density and stomatal index, the product of stomatal density and size) (Fig. 1B). Water savers are characterised by a high stomatal density, high stomatal index values, and a fast down-regulation of their conductance under increasing VPD. In contrast, water spenders down-regulate their stomatal conductance only at high VPD.

In 2019, that is, 9–10 years after planting, we selected 15 tree species to cover the trait space as well as possible by choosing species at the extremes of both gradients (two species at each end) and at intermediate values of trait expression (Fig. 1B). We used the species PCA scores on the resistance-acquisition and stomatal control trait gradient as focal tree traits, hereafter referred to as ‘focal tree resistance-acquisition traits’ and ‘focal tree stomatal control traits’. As species pools in the BEF-China experiment overlap only partly between sites A and B (Bruelheide *et al*., 2014), we sampled seven species at site A, seven at site B and one species (*Schima superba* Gardn. Et Champ.) at both sites (i.e. 15 species in total and eight species at each site) to test for potential differences between sites. We selected these species out of a pool of 25 potential tree species, which all featured mid-to fast growth rates in our experiment (Li *et al*., 2017), showed distinct radial growth rings (Böhnke *et al*., 2012), and had comparably good survival rates to ensure sufficient tree and sample size for coring.

### Focal trees and their neighbourhood

We used focal trees and their neighbours to create a realised neighbourhood species richness (NSR) gradient of 1-, 2- and 4-neighbour species (Fig. 2A,B). In the field, we randomly selected 10 focal trees (seven trees for final analysis and three trees as backup) per species (N=15) and NSR level (N=3), which resulted in 485 trees in total (one species was sampled at both sites). We sampled focal trees in as many different plots and species compositions as possible to increase the generality of our results (N=122 plots) and avoided overlapping neighbourhoods to minimise spatial-autocorrelation (see Supplementary Method 1 for details on the focal tree selection). For each focal tree, we recorded its position, species’ identity and diameter at breast height (dbh). When trees had multiple stems, the stem diameters of the two largest stems were recorded to calculate the sum of the basal areas of both stems.

**Fig. 2.**
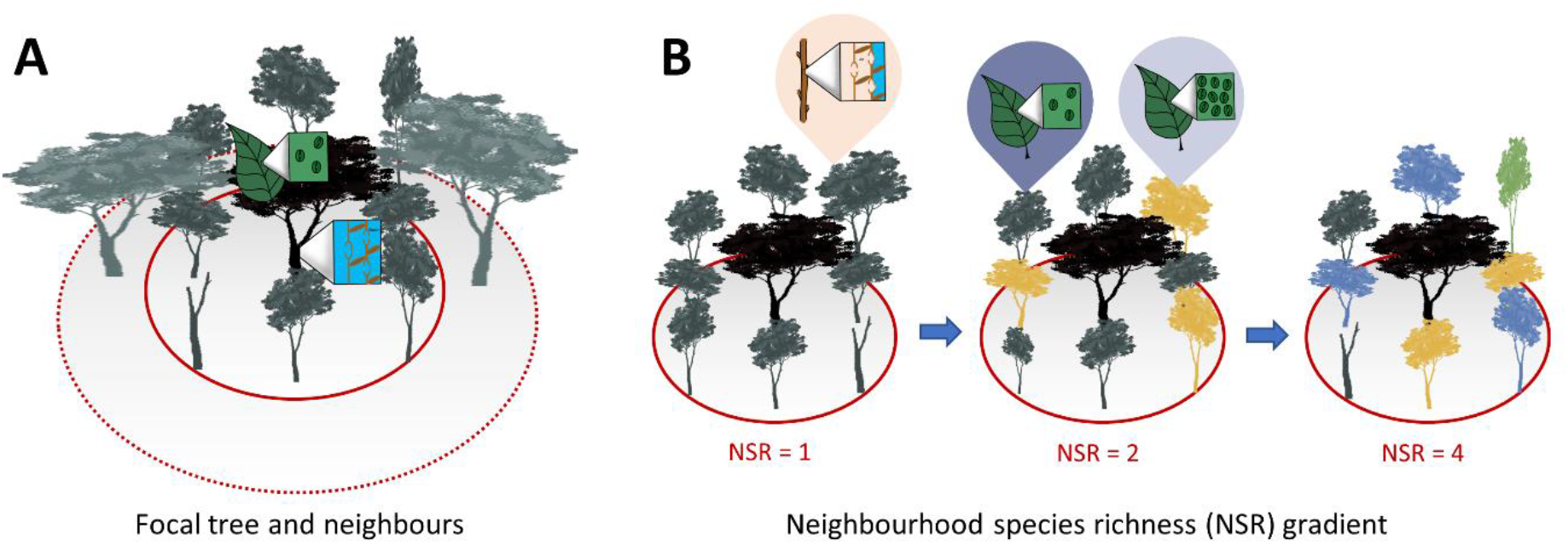
Tree neighbourhood design. **(A)** Focal trees (black tree) and their neighbours (grey trees). **(B)** Neighbourhood species richness (NSR) gradient of 1-, 2- and 4-neighbour species. The sketches illustrate the influence of focal tree and neighbour traits on the relationships between NSR, climate and growth as well as δ^13^C.

We defined a focal tree’s neighbourhood as all alive direct neighbours (occupying the immediately adjacent original planting space; maximum eight trees) and second-order neighbours (occupying the original planting spaces one further out from the direct neighbours) if their crown and the focal tree’s crown interacted (Fig. 2A). For each neighbour, we recorded its position, species’ identity, and dbh and visually estimated the height difference of neighbours compared to focal trees as a measure of shading by neighbours. We used these data to characterise the competitive environment of focal trees using eight different diameter-, height-, and distance-based neighbourhood competition indices frequently used in other studies (Table S3). We calculated neighbour traits, i.e. the functional identity of a focal trees’ neighbourhood, as the neighbourhood-weighted mean (NWM) trait value of each neighbourhood for both gradients, hereafter called ‘NWM of resistance-acquisition’ and ‘NWM of stomatal control’, similarly to the calculation of community-weighted mean traits often used in BEF studies (see, e.g. Craven *et al*., 2018) as:

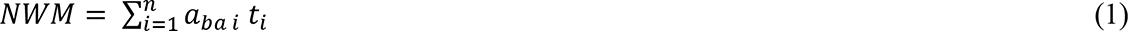

Where *a_ba_* is the abundance of species *i* measured as its basal area relative to the basal area of the other neighbour species and *t_i_* is the score of species *i* on the respective trait gradient (PCA axes reflecting resistance-acquisition or stomatal control (Fig. 1B).

### Tree growth and carbon stable isotopes

We used tree growth and carbon isotopic ratios as indicators of focal tree responses to the interactive effects of NSR, climate and drought-tolerance traits. We extracted one increment core at dbh from each focal tree perpendicular to the slope (avoiding tension wood) using a 3- threaded Haglöf increment borer with 3.5 mm core diameter. We extracted cores from the largest stem and recorded tree diameter at the coring position if coring at dbh was not possible. Cores were tightly wrapped in paper to avoid bending and dried for 72 hours at 70 °C. Core surfaces were prepared with a core-microtome (Gärtner & Nievergelt, 2010) to visualise tree-ring boundaries. Annual tree-ring width (mm) was measured using a LINTAB™ 6 system and the TSAPWin Professional 4.64 program © 2002–2009 Frank Rinn / RINNTECH with a measurement accuracy of 1/1000 mm. We measured each core twice and cross-compared series within species to ensure the correct dating of rings. No master chronology per species could be constructed owing to the short length of individual series (mostly five to seven years). Tree-ring series of 474 trees from 15 species could be dated (see Fig. S4 for an overview of wood anatomy). We used basal area increment (cm^2^) as an indicator of temporal trends in tree growth as it is, particularly in young, open-grown trees (Biondi & Qeadan, 2008), less influenced by biological age trends than tree-ring width (see Fig. S5 for a comparison). In the Supplement, we present results for tree-ring width, which yielded similar results, to allow for a comparison of both proxies following suggestions of Schwarz *et al*. (2020). Basal area increment was calculated using tree-ring width, bark thickness and diameter with the bai.out() function in the dplR package in R (Bunn *et al*., 2020).

The carbon isotopic ratio in the wood of focal trees (δ^13^C) was quantified for the years 2016- 2018 on the same cores. The rings of the years were separated, their wood homogenised and 0.8 mg woody material was weighed and placed in tin capsules. We determined δ^13^C in bulk wood rather than extracted cellulose fraction, because both materials produce highly correlated signals (Loader *et al*., 2003; Schulze *et al*., 2004). Carbon isotope analyses were conducted on an elemental analyser (NA1110, CE Instruments, Milan, Italy) coupled to a Delta+XL isotope ratio mass spectrometer (Thermo Finnigan, Bremen, Germany) via a ConFlow III at the stable isotope laboratory (BGC-IsoLab) of the Max Planck Institute for Biogeochemistry in Jena, Germany. We present carbon isotope ratio results as δ^13^C values on the VPDB-LSVEC scale (Coplen *et al*., 2006). The δ^13^C values are reported in per mil (‰) by multiplying the delta value by the factor 1000 (Coplen, 2011).

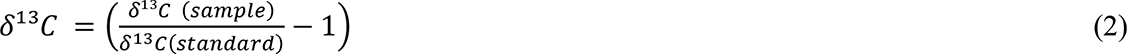

Samples were scaled against the in-house standard (acetanilide) with a δ^13^C value of -30.06 ± 0.1 ‰. Caffeine (caf-j3; δ^13^C: -40.46 ± 0.1 ‰) was analysed several times in each sequence as quality control. Linearity, blank and drift corrections were done for each sequence according to Werner & Brand (2001). We randomly remeasured a subset of samples to estimate measurement precision. The mean standard-deviation of samples from the same year and tree lay with 0.05 ± 0.03 ‰ well in the range of the in-house standard precision. We used the mean of these repeated measurements in the further analyses.

A common approach to separate the effects of water availability from those of CO_2_ assimilation on δ^13^C in forests is to analyse isotopic signals in tree rings from (co-)dominant trees that experience little shading from neighbours thereby minimizing shading effects on assimilation (Grossiord *et al*., 2014; Jucker *et al*., 2017). We selected (co-)dominant focal trees through the use of a minimum dbh of 8 cm and an *a priori* selection of those trees that experienced least shading by their neighbourhood. The final, completely balanced dataset with seven trees per species, NSR-level and year comprised 336 trees from 114 plots with at least two trees per plot.

### Statistical analysis

We used linear mixed-effects models (LMMs) to model growth (basal area increment) and wood δ^13^C responses to the interactive effects of NSR, climate and drought-tolerance traits. Our modelling framework consisted of two main steps. First, we built trait-independent neighbourhood models that accounted for the initial size of focal trees, their competition with neighbours, NSR and climate conditions, and the interaction between NSR × climate as fixed effects. We compared eight neighbourhood competition indices (Table S3) and selected the Hegyi index, which considers the distance and basal area of neighbour trees relative to the focal tree, as the best-performing one via the Akaike Information Criterion (Table S4). We accounted for our experimental design and repeated measurements through a nested random effect structure of focal tree identity nested within plot and site. A separate analysis of *Schima superba*, the species sampled at both sites, indicated that growth and δ^13^C responses did not differ between sites (Supplementary Analysis 1). Finally, we selected the most parsimonious trait-independent neighbourhood model through backward elimination of fixed and random effects. For further details see Supplementary Method 2.

Second, we examined how focal tree traits and neighbour traits modulate growth and δ^13^C responses to the interactive effects of NSR and climate. To understand the effect of focal tree traits, we included interactions between NSR, climate and either focal tree resistance-acquisition or stomatal control traits into the models of step 1. Similarly, to understand the effect of neighbour traits, we included the interaction between climate and either neighbour resistance-acquisition or stomatal control traits into the models of step 1. We selected the most parsimonious focal tree trait and neighbour trait models through backward elimination of fixed effects. Hence, in contrast to former studies (Fichtner *et al*., 2020; Jansen *et al*., 2021), we explicitly accounted for interactions between focal trees and their neighbourhood on growth and δ^13^C through modelling the effect of neighbourhood species composition not as a random effect but as a fixed effect expressed through the NWM of species’ drought-tolerance traits.

As we were interested in relative and not in absolute differences between species, we standardised basal area increment values by dividing each value by its species’ mean and δ^13^C values by subtracting its species’ mean in all mixed-effect models to reduce total variance in the data, referred to as bai_std_ and Δδ^13^C, respectively. Alternative models with species identity as a random effect yielded similar results (results not shown). LMMs were fit in R version 4.1.2 with the packages lme4 (Bates *et al*., 2015) and lmerTest (Kuznetsova *et al*., 2017). Model assumptions (normality and heteroscedasticity) were visually checked via quantile-quantile plots and through examining model residuals. We used a log transformation for basal area increment and a square-root transformation for tree-ring width to normalise residuals, centred and scaled all predictors (via subtracting µ and dividing by σ) except year and NSR before analysis and used an α of 0.05 for reporting significant effects.

## Results

Across the 15 species examined, we observed large species-specific differences in focal tree growth and particularly in δ^13^C (Fig. 3). Species responses to the interactive effects of NSR and climate were highly variable (Fig. 3). To elucidate general trends, while accounting for confounding effects such as a tree’s or a plot’s location, we employed mixed-effect models. Using trait-independent models, we found that focal tree growth, measured as basal area increment, increased logarithmically with tree size (t = 5.01, P < 0.001), decreased with competition by neighbouring trees (t = -5.91, P < 0.001), and increased by 11.4% from 1-species to 4-species neighbourhoods (t = 2.29, P = 0.024) (Fig. 4, Table S5). The effects of NSR on growth were consistent across different annual climatic conditions, and we did not observe an absolute difference in growth between the wet (2016), intermediate (2017) and dry year (2018). When examining the trait-independent models for carbon isotopic ratios (δ^13^C) in the wood of focal trees, we found a linear decrease in δ^13^C from the wet to the dry year (t = -9.06, P < 0.001; Fig. S6, Table S6). Neither tree size, competition, NSR, nor the interaction between NSR and year significantly affected δ^13^C (Tables S6,7).

**Fig. 3.**
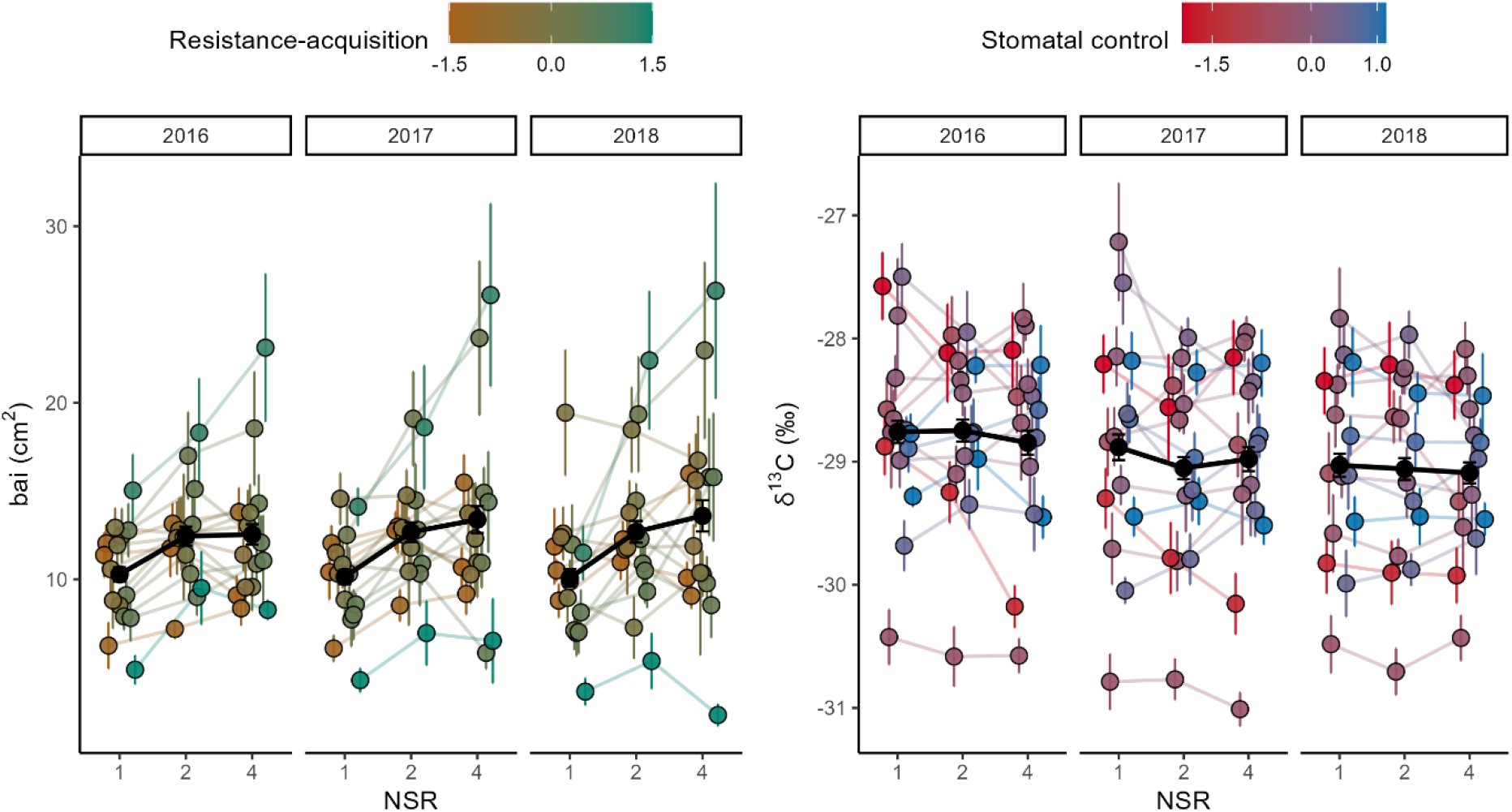
Measured values of focal tree basal area increment (bai) and carbon isotopic ratio (δ^13^C) per species, year and neighbourhood species richness (NSR) level. Points show mean values per species (N=15) and error bars respective standard errors of the mean with lines illustrating trends from 1-species to 4-species neighbourhoods. Points, error bars and lines are coloured according to the species’ trait value on either the resistance-acquisition gradient for growth or the stomatal control gradient for δ^13^C (see Fig. 1B for trait values and species identities). Black points, error bars and lines represent an overall mean and standard error of the mean across species.

**Fig. 4.**
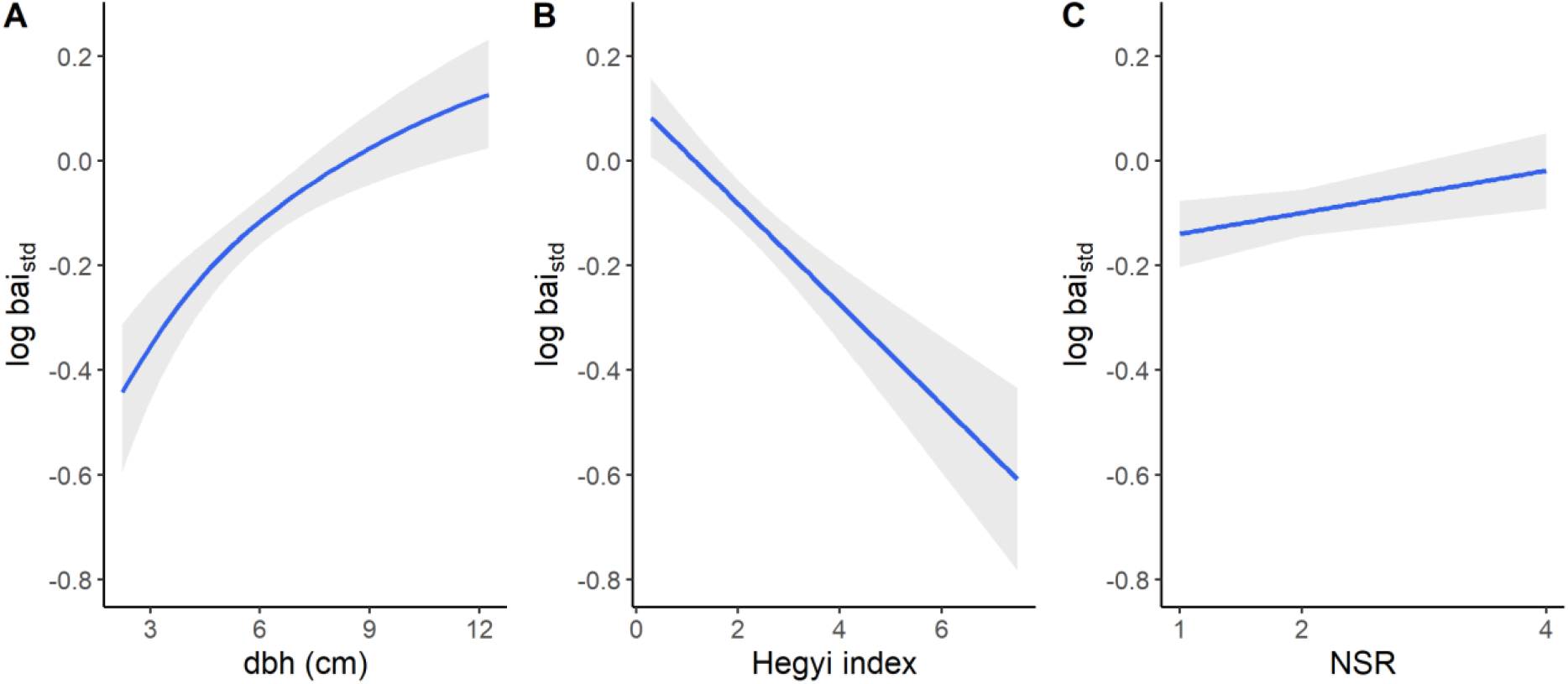
Effects of tree size (dbh), neighbourhood competition (Hegyi index) and neighbourhood species richness (NSR) on the logarithm of focal tree basal area increment (bai_std_). The blue lines are mixed-effects model fits and grey bands show a 95% confidence interval. See Table S5 for details on the fitted model.

### The effect of drought-tolerance traits on tree growth

Using trait-dependent models, we found that the resistance-acquisition traits of focal trees significantly modulated the relationship between tree growth and two factors: NSR and climate (Fig. 5A,B, Table 1). With increasing NSR, acquisitive species grew 29.7 % more in 4-species compared to 1-species neighbourhoods. On the other hand, resistant species experienced a 8.9% decrease in growth in 4-species compared to 1-species neighbourhoods (NSR × focal tree resistance-acquisition traits, t = 2.45, P = 0.015; Fig. 5A). Furthermore, from the wet to the dry year, growth increased for resistant species but declined for acquisitive species (year × focal tree resistance-acquisition traits, t = -6.84, P < 0.001; Fig. 5B). We also found that resistance-acquisition traits of focal trees had contrasting effects on the relationship between NSR, climate, and growth, although this three-way interaction was only marginally significant (NSR × year × focal tree resistance-acquisition traits, t = -1.75, P = 0.080; Fig. S7, Table S16): In the wet year, acquisitive species benefited from higher NSR, while resistant species grew less. In the dry year, the effects were weaker but predominately positive, with acquisitive species still growing more with higher NSR, while resistant species were unaffected by NSR. Notably, the resistance-acquisition traits of neighbours significantly influenced growth responses. Focal trees in a neighbourhood dominated by resistant species grew more from the wet to the dry year, while those in an acquisitive neighbourhood grew less (year × NWM of resistance-acquisition, t = - 4.17, P < 0.001; Fig. 5C, Table 1).

**Fig. 5.**
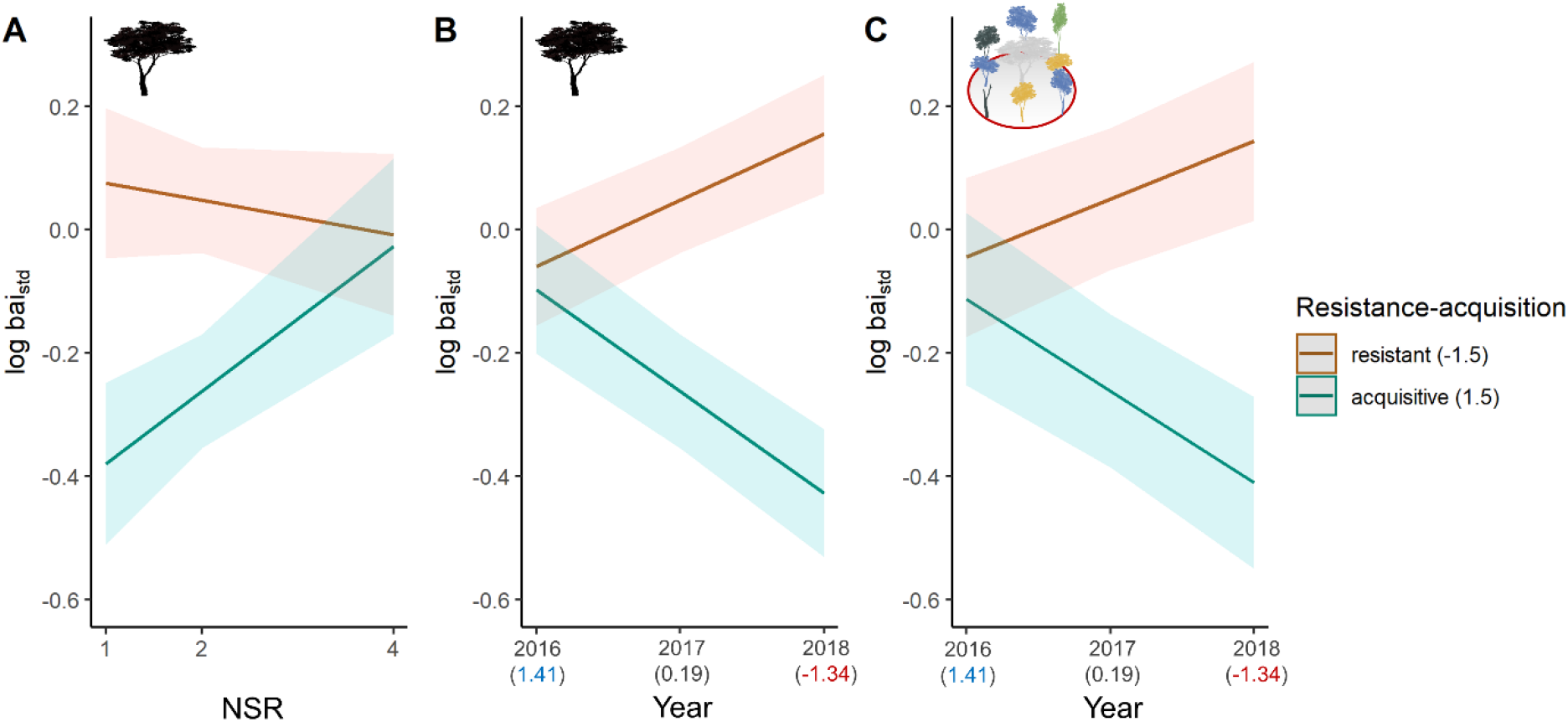
Modulation of the relationship between neighbourhood species richness (NSR) and growth and between climate and growth by resistance-acquisition traits. Lines represent linear mixed-effects model fits and coloured bands show a 95% confidence interval. The models depict significant effects of NSR and study year (2016-2018 with wet to dry climate, SPEI values in brackets) on the logarithm of basal area increment (bai_std_) of focal trees predicted for resistant (PC1 value of -1.5) and acquisitive species (PC1 value of 1.5). The panels illustrate the influence of focal tree resistance-acquisition traits (black tree; A, B) and neighbour resistance-acquisition traits (C) on the relationships. See Fig. 1-2 for details on the study design and Table 1 for the fitted models.

**Table 1.** Mixed-effect model statistics for effects of tree size, competition by neighbours, neighbourhood species richness (NSR), year and drought-tolerance traits and interactions on the basal area increment and δ^13^C of focal trees. Models were fit for resistance-acquisition or stomatal control traits using either focal tree traits or neighbour traits; individual models are separated by grey shading. t and F indicate t and F statistics based on Satterthwaite’s method and p the p-value of the significance test.

| Source of variation | Resistance-acquisition |  |  |  | Stomatal control |  |  |  |
| --- | --- | --- | --- | --- | --- | --- | --- | --- |
|  | Focal tree traits |  | Neighbour traits |  | Focal tree traits |  | Neighbour traits |  |
|  | t | p | t | p | t | p | t | p |
| **Basal area increment ($\text{bai}_{\text{std}}$)** |  |  |  |  |  |  |  |  |
| Size | 5.8 | <0.001 | 5.6 | <0.001 | 5.0 | <0.001 | 5.0 | <0.001 |
| Competition | -6.1 | <0.001 | -6.1 | <0.001 | -5.9 | <0.001 | -5.9 | <0.001 |
| NSR | 2.4 | 0.018 | 2.1 | 0.036 | 2.3 | 0.024 | 2.2 | 0.027 |
| Year | -1.8 | 0.074 | -1.8 | 0.080 | -1.8 | 0.078 | -1.7 | 0.082 |
| Trait | -0.3 | 0.758 | 1.1 | 0.282 | 3.4 | 0.001 | 2.6 | 0.011 |
| NSR × Year | - | - | | | - | - | | |
| NSR × Trait | 2.5 | 0.015 | | | - | - | | |
| Year × Trait | -6.8 | <0.001 | -4.2 | <0.001 | -5.1 | <0.001 | -3.0 | 0.002 |
| NSR × Year × Trait | - | - | | | - | - | | |
| **Carbon isotopic ratio ($\Delta\delta^{13}\text{C}$)** |  |  |  |  |  |  |  |  |
|  | F | p | F | p | t | p | t | p |
| Size | - | - | - | - | - | - | - | - |
| Competition | - | - | - | - | - | - | - | - |
| NSR | - | - | - | - | -0.9 | 0.351 | - | - |
| Year | 43.6 | <0.001 | 44.0 | <0.001 | -4.8 | <0.001 | -9.1 | <0.001 |
| Trait | 0.0 | 0.847 | 0.2 | 0.649 | -2.8 | 0.005 | -2.2 | 0.028 |
| NSR × Year | - | - | | | 0.5 | 0.610 | | |
| NSR × Trait | - | - | | | 2.3 | 0.024 | | |
| Year × Trait | 6.2 | 0.002 | 9.4 | <0.001 | 3.7 | <0.001 | 3.4 | 0.001 |
| NSR × Year × Trait | - | - | | | -2.7 | 0.008 | | |

Note: Only the most parsimonious models are presented; parameters dropped during model selection are indicated with '-'. See Fig. 1-2 for details on the study design and Tables S8-S15 for details on the fitted models.

The stomatal control traits of focal trees significantly modulated the relationship between climate and growth, but not the relationship between NSR and growth nor between NSR, climate and growth (Fig. 6A, Table 1). Specifically, water-saving species increased in their growth, while water-spending species decreased in their growth from the wet to the dry year (year × focal tree stomatal control traits, t = -5.10, P < 0.001; Fig. 6A). Furthermore, the stomatal control traits of neighbours significantly influenced the climate-growth relationship: Focal trees in a neighbourhood dominated by water-saving species grew better from the wet to the dry year, while growth of focal trees in a water spending neighbourhood declined (year × NWM of stomatal control, t = -3.04, P = 0.002; Fig. 6B, Table 1). The growth responses, as found for basal area increment, were similar for tree-ring width, except for a general decline in tree-ring width from 2016-2018 and with tree size, presumably due to an age trend (Figs. S8- 10). For tree-ring width, the three-way interaction between NSR, year and the resistance-acquisition traits of focal trees was significant (t = -2.21, P = 0.027; Fig. S9).

**Fig. 6.**
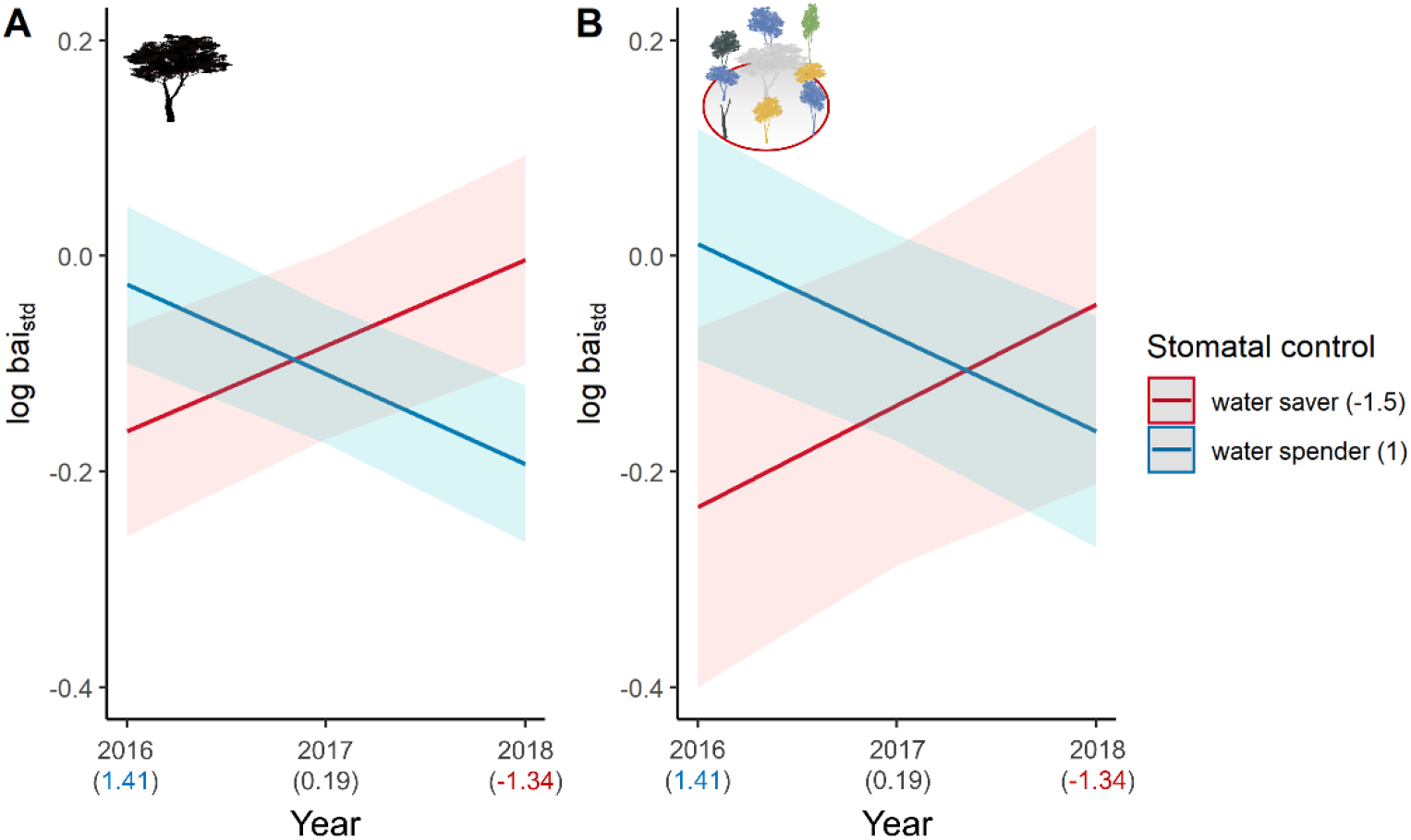
Modulation of the relationship between climate and growth by stomatal control traits. Lines represent linear mixed-effects model fits and coloured bands show a 95% confidence interval. The models depict significant effects of study year (2016-2018 with wet to dry climate, SPEI values in brackets) on the logarithm of basal area increment (bai_std_) of focal trees predicted for water savers (PC2 value of -1.5) and water spenders (PC2 value of 1.0). The panels illustrate the influence of focal tree stomatal control traits (black tree; A) and neighbour stomatal control traits (tree neighbourhood; B) on the relationship. See Fig. 1-2 for details on the study design and Table 1 for the fitted models.

### The effect of drought-tolerance traits on δ^13^C

Using trait-dependent models to explain variations in δ^13^C in the wood of focal trees, we found significant interactions of resistance-acquisition traits with climate, with a stronger influence observed from neighbour traits compared to focal tree traits (Fig. 7, Table 1). In both the intermediate and dry year, we observed a tendency towards higher δ^13^C (indicating higher water stress) in focal trees located in acquisitive neighbourhoods compared to resistant neighbourhoods (Fig. 7B). However, in the wet year, lower δ^13^C (indicating lower water stress) occurred in focal trees situated in acquisitive neighbourhoods (year × NWM of resistance-acquisition, year as a categorical fixed effect, F = 9.45, P < 0.001; Fig. 7B). The difference between focal trees in resistant *vs* acquisitive neighbourhoods was most prominent during the year with intermediate water availability (2017; Fig. 7B).

**Fig. 7.**
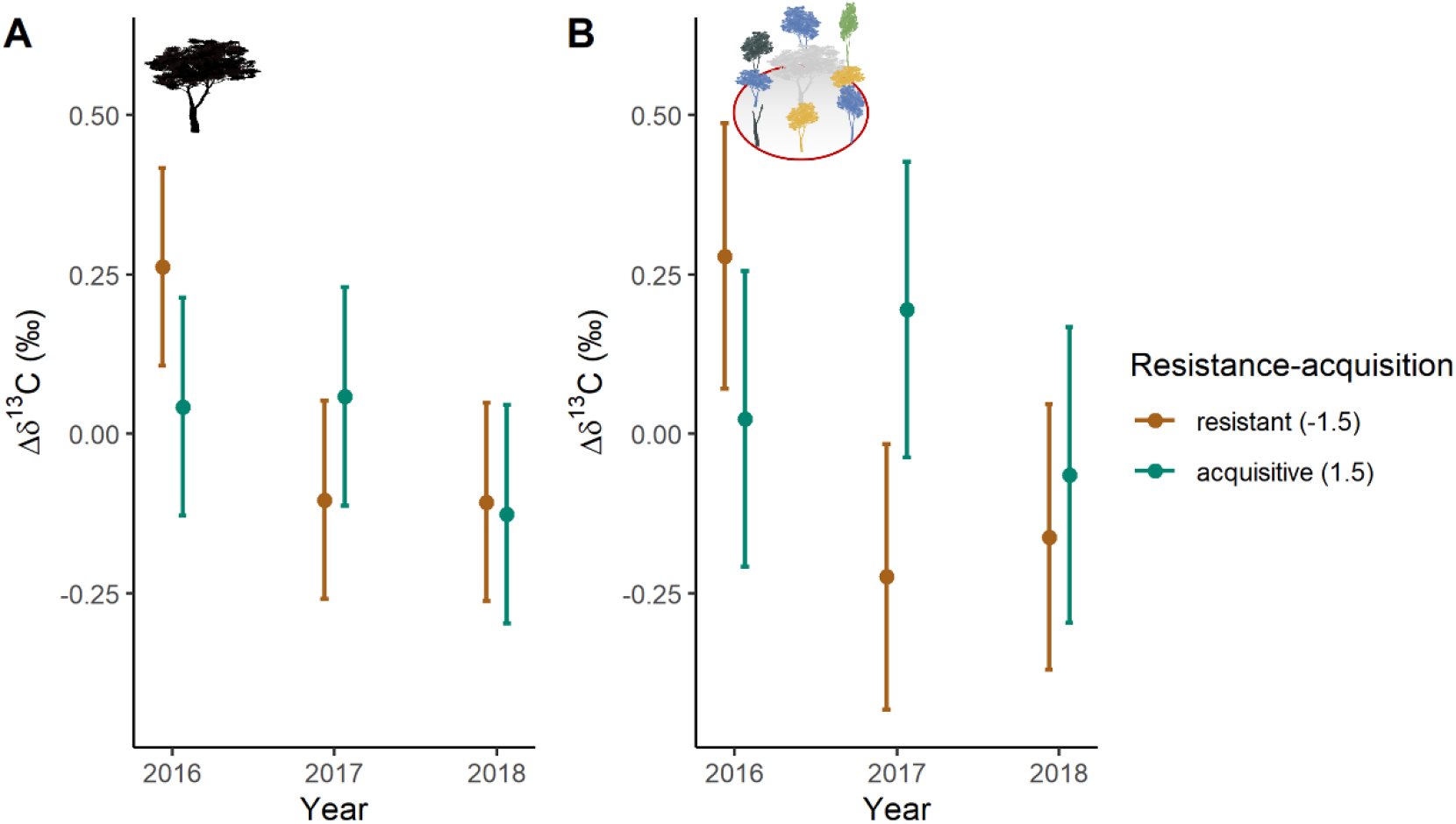
Modulation of the relationship between climate and δ^13^C by resistance-acquisition traits. Points represent linear mixed-effects model fits and error bars show a 95% confidence interval. The models depict significant effects of study year (2016-2018 with wet to dry climate, SPEI values in brackets) on δ^13^C in the wood of focal trees predicted for resistant (PC1 value of -1.5) and acquisitive species (PC1 value of 1.5). The panels illustrate the influence of focal tree resistance-acquisition traits (black tree; A) and neighbour resistance-acquisition traits (tree neighbourhood; B) on the relationship. See Fig. 1-2 for details on the study design and Table 1 for the fitted models.

The stomatal control traits of focal trees played a significant role in modulating the relationship between NSR, climate and δ^13^C (NSR × year × focal tree stomatal control traits, t = -2.66, P = 0.008; Fig. 8, Table 1): We found contrasting NSR effects on δ^13^C for water-saving and water-spending species, which weakened as climate conditions transitioned from wet to dry (Fig. 8). In the wet year, water-saving species exhibited a decrease in δ^13^C with increasing NSR. However, this positive diversity effect declined in the dry year, resulting in similar δ^13^C across NSR levels. In contrast, water-spending species tended to show an increase in δ^13^C with increasing NSR in the wet year, but a decrease in δ^13^C with increasing NSR in the dry year. Hence, water-saving species benefited from more species-rich neighbourhoods (NSR) by experiencing lower water stress during wet conditions. In contrast, water-spending species tended to benefit from higher NSR during dry climatic conditions, although the effect sizes were relatively small. Finally, the stomatal control traits of neighbours influenced the δ^13^C in the wood of focal trees (year × NWM of stomatal control, t = 3.43, P = 0.001; Fig. S11, Table 1). Focal trees surrounded by water-saving neighbours had lower δ^13^C in the dry year compared to the wet year, but this effect may have been enhanced by the overall decline in δ^13^C from the wet to the dry year (Fig. S6).

**Fig. 8.**
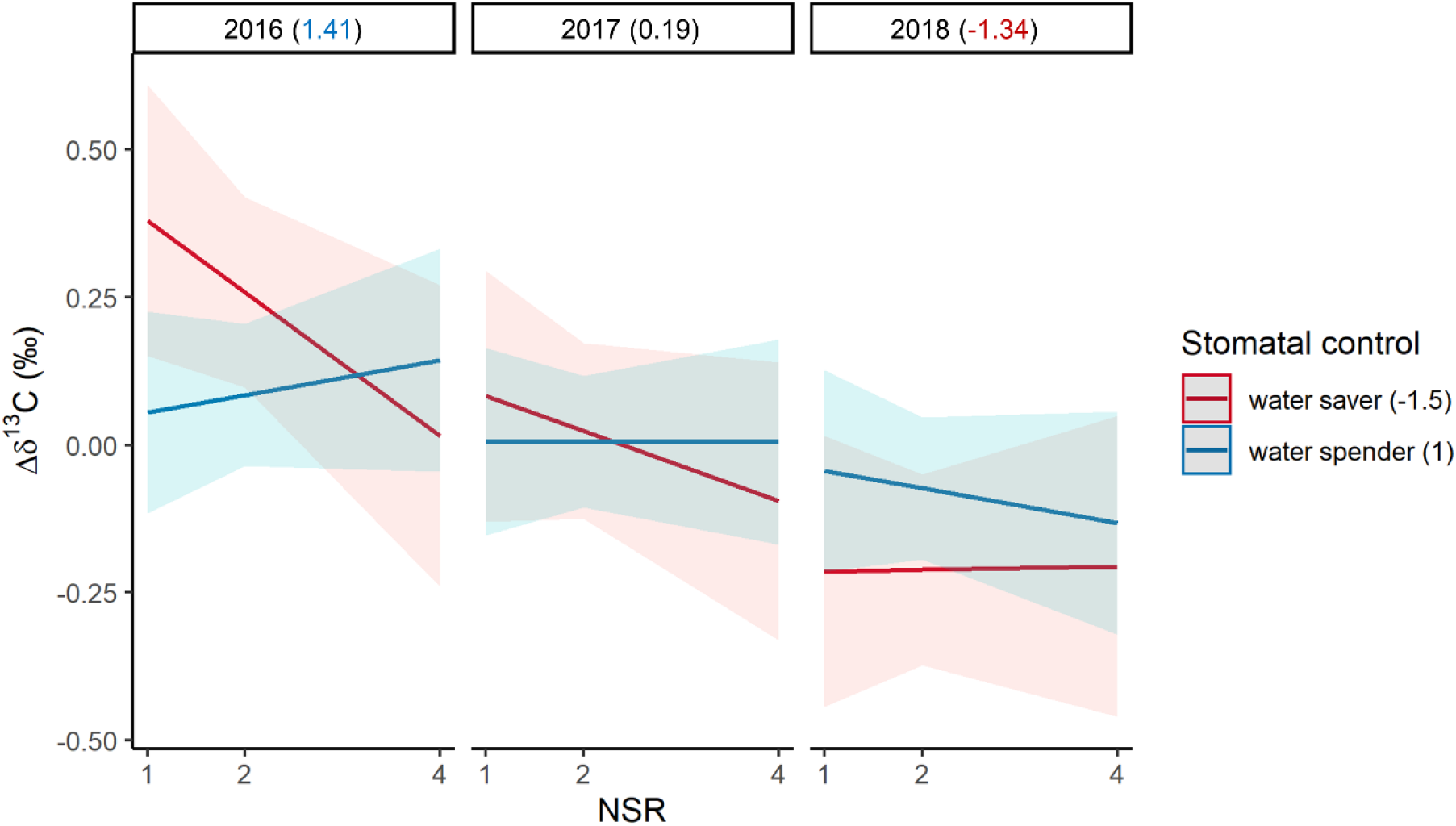
Modulation of the relationship between neighbourhood species richness (NSR), climate and δ^13^C by stomatal control traits. Lines represent linear mixed-effects model fits and coloured bands show a 95% confidence interval. The models depict significant, interactive effects of NSR and study year (2016- 2018 with wet to dry climate, SPEI values in brackets) on δ^13^C in the wood of focal trees predicted for water savers (PC2 value of -1.5) and water spenders (PC2 value of 1.0). See Fig. 1-2 for details on the study design and Table 1 for the fitted model.

## Discussion

The growth of focal trees increased with neighbourhood tree species richness (NSR) across the 15 species examined. However, we did not find an overall significant effect of NSR on carbon isotopic ratios in the wood of focal trees (δ^13^C), which we considered as a proxy of neighbourhood scale water availability (Grossiord *et al*., 2014; Jucker *et al*., 2017). Moreover, the strength of the relationship between NSR and growth and between NSR and δ^13^C did not increase from wet to dry climatic conditions. Instead, relationships between NSR, climate and growth or δ^13^C were modulated by drought-tolerance traits of focal trees and their neighbours regarding cavitation resistance vs. resource acquisition and stomatal control. The biotic context (i.e. focal tree and neighbour traits) thus determined the strength and nature of BEF relationships in the examined tree communities. Resistance-acquisition traits primarily modulated NSR and climate effects on growth, whereas stomatal control traits primarily influenced wood δ^13^C. As expected, growth was thus more strongly related to the leaf economics spectrum (Reich, 2014), while δ^13^C was controlled by stomatal aperture (Farquhar *et al*., 1989).

### Tree diversity increases growth but does not universally relieve drought stress

As postulated in H1, NSR increased growth, which is consistent with findings from other studies (Guillemot *et al*., 2020; Schnabel *et al*., 2019; Trogisch *et al*., 2021), including those from our experiment (Fichtner *et al*., 2018; Fichtner *et al*., 2020). Still, NSR effects on the growth of individual species can be both positive and negative and vary with climatic conditions (Fig. 3; e.g. Vitali *et al*., 2018). Tree growth is an integrated signal of many biotic and abiotic drivers (Grossiord, 2020). The positive effect of NSR on growth is thus likely the result of different and interacting mechanisms operating at the neighbourhood scale (Trogisch *et al*., 2021). The positive interactions potentially include resource partitioning (of light, water and nutrients), abiotic facilitation (such as microclimate amelioration) and biotic interactions (such as dilution of generalist pathogens) (Barry *et al*., 2019).

Contrary to our expectation (H1) and an earlier study on twig δ^13^C in our experiment conducted during the particularly wet year 2015 (Fig. 1A; Jansen *et al*., 2021), we did not detect a general enhancement of water availability in more diverse tree neighbourhoods. Similarly, other studies found mixed results and no universal decrease in δ^13^C in mixtures compared to monocultures (Grossiord, 2020; Haberstroh & Werner, 2022). The net negative effect of NSR on δ^13^C in Jansen *et al*. (2021) is likely due to increased shading at high NSR, which reduces photosynthesis rates, and not due to enhanced water availability. Our assumption is based on the fact that Jansen *et al*. (2021) found the strongest NSR effects for trees with a high degree of crown competition from neighbours. In contrast, our δ^13^C values should be primarily influenced by NSR-and climate-induced changes in neighbourhood-scale water availability as we only studied (co-)dominant trees with slight shading. This notion is supported by the non-significant effect of competition indices on δ^13^C. Moreover, drought affects stomatal conductance more than CO_2_ assimilation (Farquhar *et al*., 1989; Grossiord *et al*., 2014), supporting the importance of water availability as main driver of δ^13^C values in our study.

To understand the abiotic context dependency of BEF relationships, we selected years to represent a gradient from wet to dry climatic conditions (SPEIs; Fig. S1). Contrary to our expectation (H1), we did not observe a consistent increase in diversity effects with increasing dryness (SPEIs; Fig. S1), suggesting that water was not limiting (even though the examined dry year had similar SPEIs as past drought years; Fig. S2) or that drought impacts were buffered through the mobilization of reserves (McDowell *et al*., 2022). Alternatively, light may have been the primary limiting factor (Forrester & Bauhus, 2016). These points may also explain why we observed no overall decline in tree growth and decreasing δ^13^C values from the wet to the dry year. Hence, even though all forest biomes, including the comparably humid subtropical forests examined here, are threatened by drought (Hartmann *et al*., 2022), responses may be more pronounced during extreme droughts (e.g. during hotter droughts; Schnabel *et al*., 2022).

Other studies reported stronger diversity effects during dry (Fichtner *et al*., 2020; Schnabel *et al*., 2019) and during wet years (Belluau *et al*., 2021). Similarly, positive, neutral and negative effects of species richness on growth and δ^13^C have been reported under drought (Grossiord, 2020). An important difference between these former and our study is that we examined a balanced trait sample (in terms of species and number of individuals) along both drought-tolerance trait gradients (Fig. 1B). Given that species with contrasting drought-tolerance traits showed opposite responses to NSR and climate in our study, it is not surprising that we detected a net zero or only a weak overall effect of NSR on focal tree responses to drought. In contrast, the non-balanced trait sample in former studies may explain their mixed results on the modulation of drought impacts on trees and forests by tree diversity. Hence, the contrasting water-use strategies of focal trees and their neighbours may be one fundamental missing link for understanding the role of tree diversity during drought. Here, we attempted to address this by considering the biotic context dependency of BEF relationships, which we quantified in terms of their modulation by drought-tolerance traits of focal trees and their neighbours.

### The importance of focal tree traits

Consistent with H2, we found that during drought, NSR increased growth and decreased wood δ^13^C of acquisitive and water-spending focal tree species. In contrast, resistant and water-saving species did not respond to NSR, presumably because species interactions did not aid their strategies to cope with drought. Hence, high neighbourhood diversity supported the more vulnerable species in the forest community during drought (Fichtner *et al*., 2020). In general, the presence of heterospecific neighbours is more likely with increasing NSR. These heterospecific neighbours can influence focal tree responses by enhancing or reducing competition or promoting facilitation (Forrester, 2017; Forrester & Pretzsch, 2015). In our study, acquisitive species likely grew better due to their ability to acquire and use resources and the lower inter-compared to intraspecific competition for water at high NSR (Fichtner *et al*., 2017; Fichtner *et al*., 2020). Similarly, NSR may have relieved water stress for water-spending species during drought by increasing the likelihood of having water-saving neighbours, which transpire less water, thereby increasing local soil water availability (Forrester, 2017). The lower physiological water stress (lower δ^13^C) we found in water-spending species during drought at high NSR indicates a novel drought mitigation effect of diversity in addition to the protection of acquisitive species reported formerly (Fichtner *et al*., 2020). The growth decline we observed for resistant and water-saving species from the wet to the dry year could be related to higher degrees of embolism in the vessels of acquisitive species (which are sensitive to cavitation) and higher risks for such cavitation in water-spending species (McDowell *et al*., 2008). In contrast, their less vulnerable counterparts may have been able to grow better under drought.

### The importance of neighbour traits

Consistent with H3, we found that the drought-tolerance traits of tree neighbours consistently changed the nature of focal tree growth and δ^13^C responses to climatic conditions. These changes by the traits of neighbours operated in the same direction as the changes induced by the traits of focal trees, thereby amplifying tree responses to climate. As expected, focal trees grew less in neighbourhoods dominated by acquisitive and water-spending species during drought, while they grew more in resistant and water-saving neighbourhoods. We also observed higher δ^13^C values in focal trees in acquisitive neighbourhoods during the intermediate-and dry-year. Both observations indicate reductions in local water availability and thus enhanced water stress (Forrester, 2017; Grossiord *et al*., 2014) in neighbourhoods dominated by acquisitive and water-spending species relative to neighbourhoods dominated by resistant and water-saving species. This finding may be explained by acquisitive and water-spending neighbours having a higher water consumption during drought. Acquisitive species tend to have a high maximum xylem hydraulic conductance (Bongers *et al*., 2021), while water spenders close their stomata only late during dry conditions and thus continue to transpire and consume water (McDowell *et al*., 2008).

Fichtner *et al*. (2020) already suggested that reduced competition for water between heterospecific neighbours benefits cavitation-sensitive species in diverse neighbourhoods. Still, they could not test this assumption as they did not quantify the influence of neighbour traits. Explicitly accounting for neighbour traits – instead of using neighbourhood species composition as a random effect (e.g. Fichtner *et al*., 2020; Jansen *et al*., 2021) – allowed us to quantify the influence of the functional identity of neighbourhoods and the influence of their diversity.

### Outlook

Trait-based neighbourhood models have been used to understand tree growth, survival and community assembly (Kunstler *et al*., 2012; Uriarte *et al*., 2010; Yang *et al*., 2021) and in few cases to understand how focal tree traits modulate tree growth responses to species richness and climate (e.g. Fichtner *et al*., 2020). However, despite compelling arguments for why the traits of neighbouring trees may alter local water availability (Forrester, 2017), we accounted here for the first time for the influence of focal tree and neighbour traits on BEF relationships in experimental tree communities. The orthogonality of the resistance-acquisition and stomatal control gradient in our study system (Fig. 1B) allowed us to disentangle the respective contributions of both gradients through exploring the effects of stomatal control (or resistance-acquisition) traits at mean levels of the other gradient. The species responses in years with different water availability were consistent with the current understanding of trade-offs between high cavitation resistance (low Ψ_50_) and acquisitive resource use (Guillemot *et al*., 2022; Reich, 2014) and between water-spending and water-saving stomatal control (Martínez-Vilalta & Garcia-Forner, 2017). We discuss this and potential relationships between resistance-acquisition and stomatal control traits in detail in Supplementary Discussion 1. However, despite our consideration of multiple traits, future studies could foster a more holistic understanding of tree drought responses and their modulation by functional traits through studying the role of traits not considered here such as the storage of non-structural carbohydrates (Hajek *et al*., 2022; McDowell *et al*., 2022) and attributes of the root system (Weigelt *et al*., 2021). Moreover, in cases where both focal tree and neighbour traits influenced observed responses, our modelling approach does not allow to mechanistically disentangle their relative contributions. Future studies could improve our ability to do so by sampling (or planting) trees along orthogonal gradients in focal tree and neighbour traits. Finally, despite the rapid tree growth in the subtropics that enabled us to study fairly large trees (25% of the experimental communities had grown taller than 10m; Schnabel *et al*., 2021), the juvenile growth trajectories may not have revealed trends that become apparent only at an older age.

Overall, our results extend research on the modulation of drought impacts on forests by tree diversity, beyond observational studies in comparably species poor forests (Grossiord, 2020) to include species-rich tree communities featuring contrasting drought-tolerance traits which are grown under experimental conditions. This approach enabled the establishment of causality while minimizing confounding factors such as environmental variation. We could derive two key conclusions: (1) Considering the functional identity of focal trees and their neighbours resolved the biotic context dependency of BEF relationships; such a trait-based perspective may help to explain positive and negative mixing effects under drought. (2) Drought-tolerance traits of focal trees and interactions with their tree neighbours induced contrasting species responses to wet *vs* dry climatic conditions; this trait-driven species asynchrony is a key driver of positive diversity-stability relationships in forests (Schnabel *et al*., 2021). The biotic context we analysed using a trait-based approach, with traits tailored to the target ecosystem functions assessed, may help to generalize the context dependency of BEF relationships and is relevant for designing tree mixtures suitable to cope with a range of different drought conditions. It can give insight into the optimal configuration of tree neighbourhoods in terms of diversity and identity in drought-tolerance traits. This may help to optimise forest productivity and foster stability to climate extremes.

## Acknowledgements

We thank local workers, in particular Mr. Wang and Mr. Shi, for invaluable help in the field, Luise Münsterberg, Paulina Tjandraputri and Lara Schmitt for supporting the sample preparation for the carbon isotope analyses and Wenzel Kröber and Helge Bruelheide for trait measurements. This research was supported by the International Research Training Group TreeDì funded by the Deutsche Forschungsgemeinschaft (DFG, German Research Foundation) – 319936945/GRK2324 and the University of Chinese Academy of Sciences.

## Author contributions

Florian Schnabel, Kathryn E. Barry, Anja Kahl, Joannès Guillemot, Jürgen Bauhus and Christian Wirth conceived the ideas of the study and designed the methodology; Florian Schnabel, Susanne Eckhardt, Heike Geilmann, Anja Kahl and Heiko Moossen collected the data; Florian Schnabel and Susanne Eckhardt analysed the data and Kathryn E. Barry, Joannès Guillemot, Jürgen Bauhus and Christian Wirth joined the interpretation of the data and results; Florian Schnabel created the figures; Florian Schnabel wrote the manuscript; All authors contributed critically to the drafts and gave final approval for publication.

## Data availability statement

Tree growth and stable carbon isotope data are available from the BEF-China project database (https://data.botanik.uni-halle.de/bef-china; data will be made publicly available before publication). Climate and trait data are available from the Climatic Research Unit (CRU TS v4.04; Harris *et al*., 2020) and from Kröber *et al*. (2014), respectively.

