## Supplementary Material for "Neighbourhood species richness and drought-tolerance traits modulate tree growth and δ^13^C responses to drought"

**This file includes:**

- Figs. S1 to S11
- Tables S1 to S18
- Supplementary Methods 1 – 2
- Supplementary Analysis 1
- Supplementary Discussion 1
- References

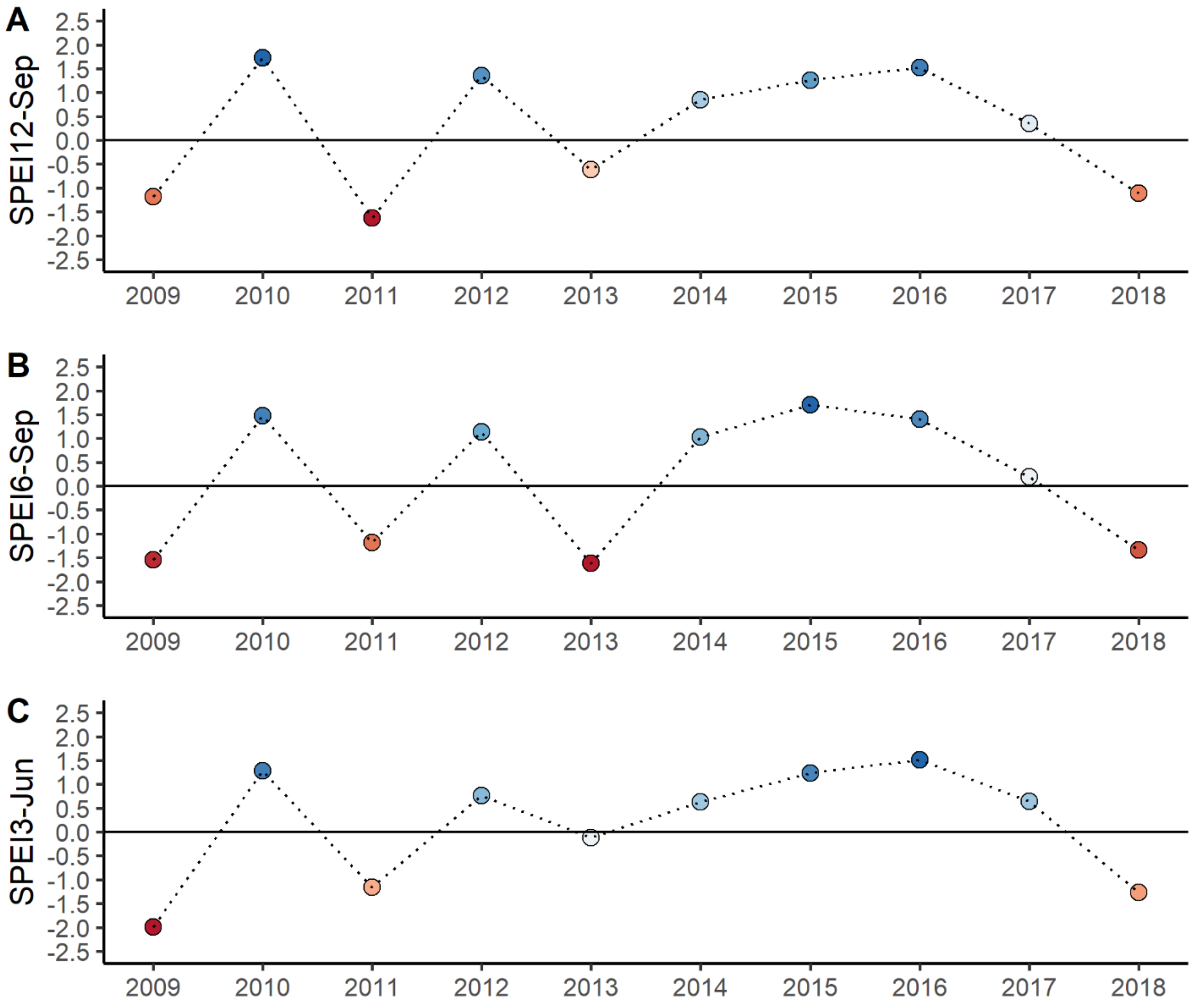


**Fig. S1** Climate-based characterisation of the study years 2016 (wet), 2017 (intermediate) and 2018 (dry). Shown are standardised climatic water balances calculated based on the standardised precipitation evapotranspiration index (SPEI) (Vicente-Serrano *et al.*, 2010) calculated from a high-resolution time-series of interpolated climate station data (CRU TS v4.04; Harris *et al.*, 2020). SPEIs are compared for the three months of the peak vegetation period (SPEI3, April-July) for the six months of the entire vegetation period (SPEI6, April-September) and the twelve months of a whole year since the end of the vegetation period of the preceding year (SPEI12, October-September), since the establishment of the BEF-China experiment (2009). The wet-to-dry study years are highlighted with a red circle. Blue points indicate wetter and red points drier conditions than the long-term mean (1901-2019); values below -1 and above 1 can be considered exceptional.

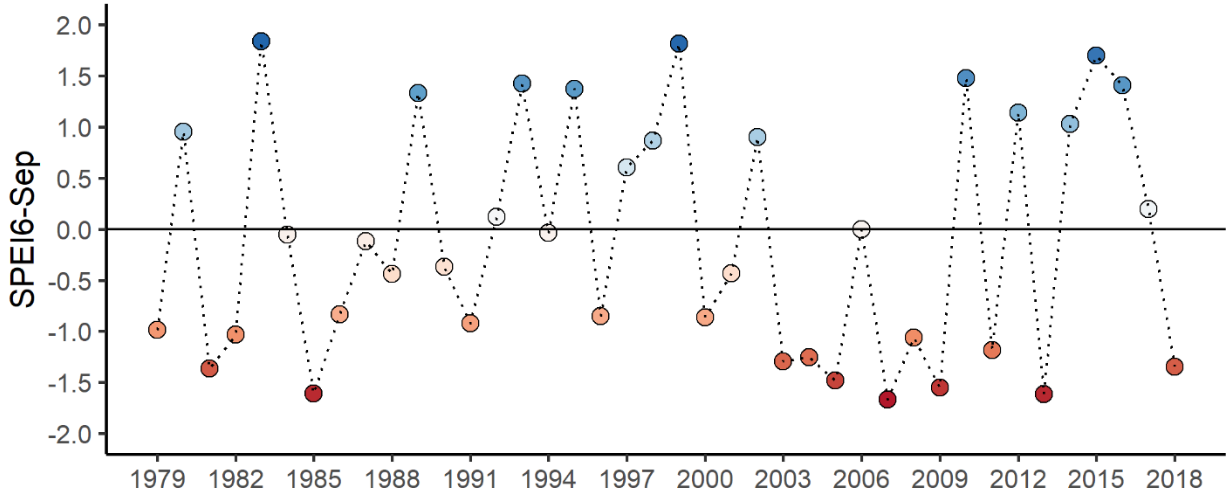


**Fig. S2** A long-term perspective on standardized climatic water balances at our study site. Shown are values of the standardised precipitation evapotranspiration index (SPEI) for the principal vegetation period (April-September). For further details on the underlying data and SPEI calculation see methods and Fig.S1.

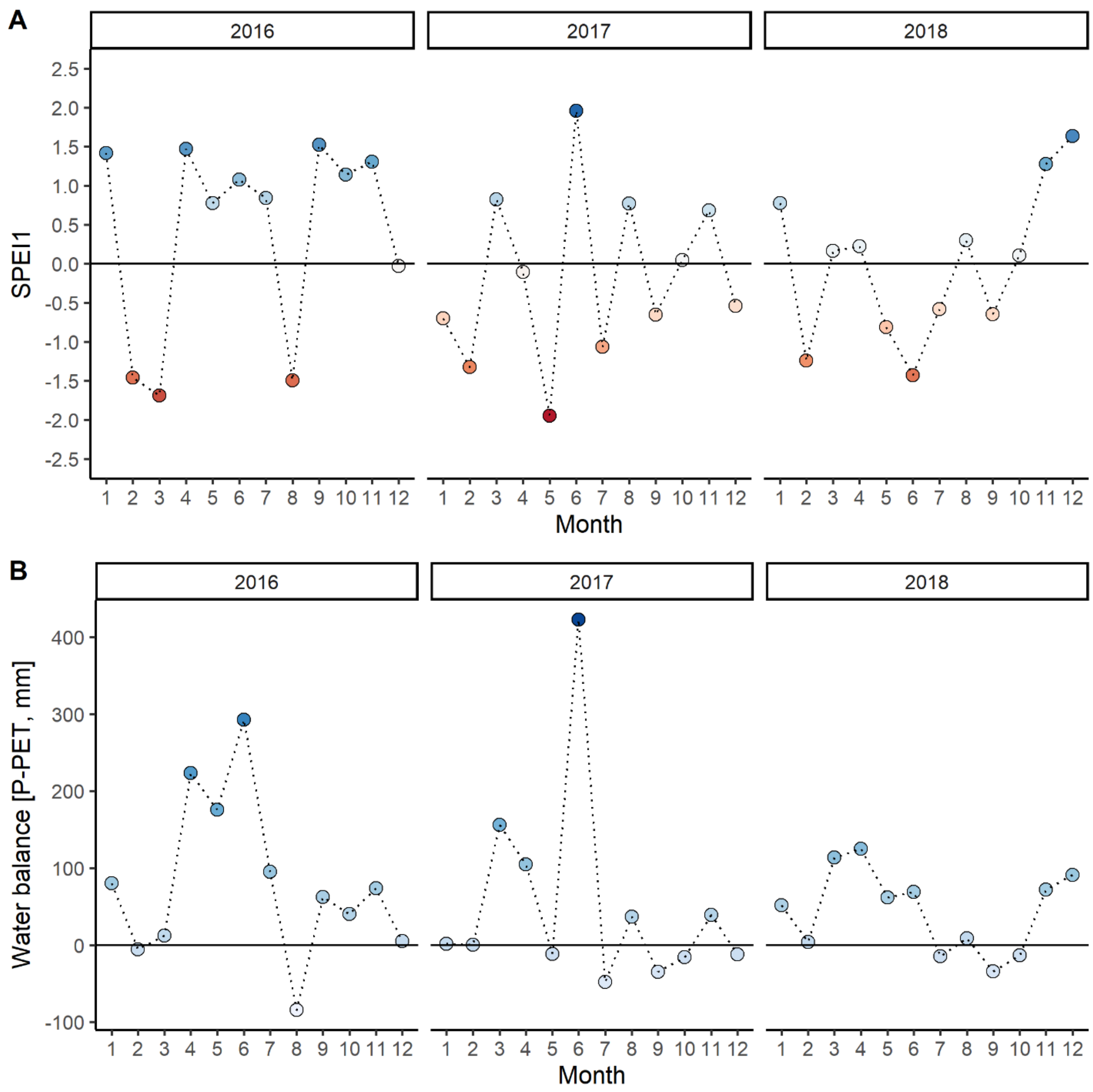


**Fig. S3** Intra-annual climatic water balances at our study site. Shown are (A) values of the standardised precipitation evapotranspiration index (SPEI) and (B) non-standardized water balances of precipitation minus potential evapotranspiration (PET) for each month for the study years 2016-2018. For further details on the underlying data and SPEI calculation see methods and Fig.S1.

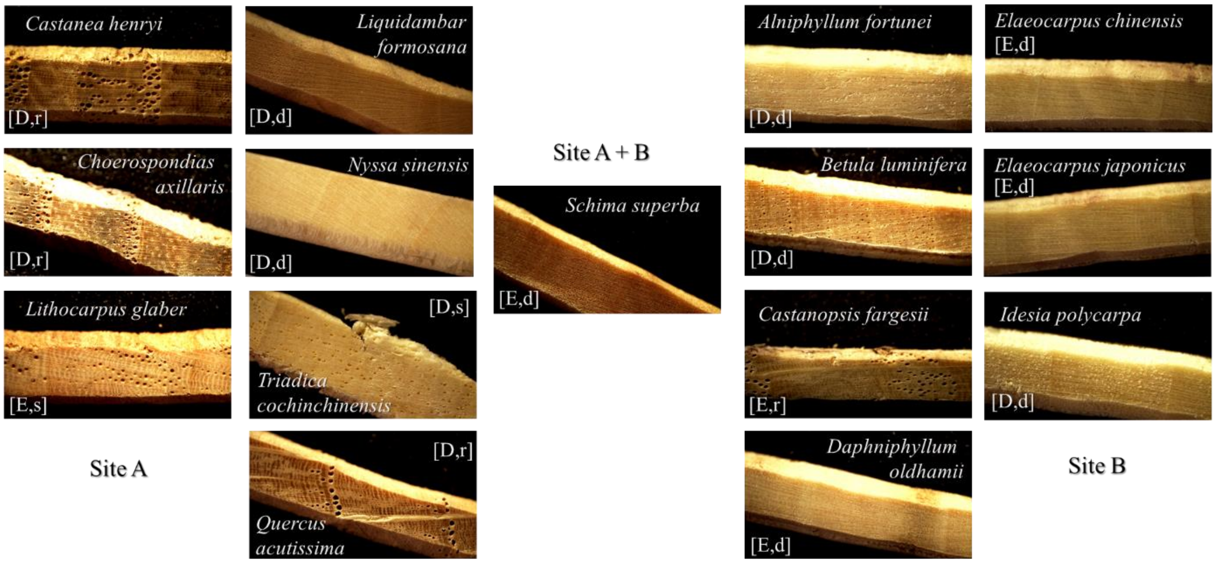


**Fig. S4** Wood anatomy of the 15 tree species sampled in this study. Shown are photographs of exemplary cores per species and experimental site (see Table S1 for details on the species). Capital letters show the leaf habit (evergreen (E) and deciduous (D)) and lower-case letters wood porosity (ring porous (r), diffuse porous (d) and semi-ring porous (s)). Photographs were taken after surface preparation with a core microtome (Gärtner & Nievergelt, 2010).

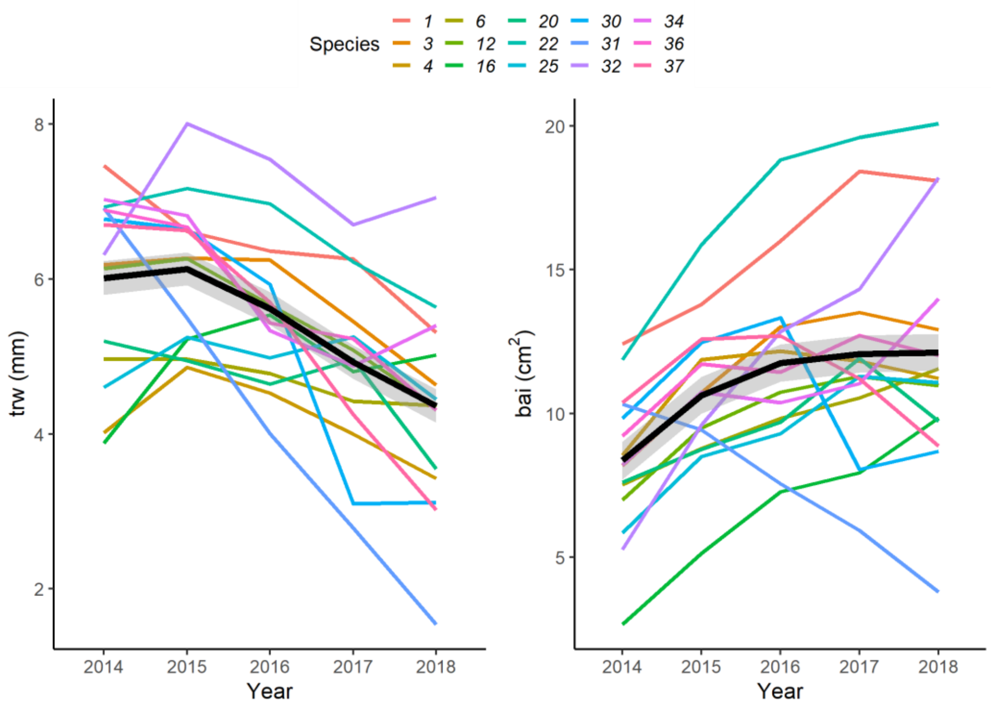


**Fig. S5** Comparison of focal tree tree-ring width (trw, mm) and basal area increment (bai, $cm^2$) series.

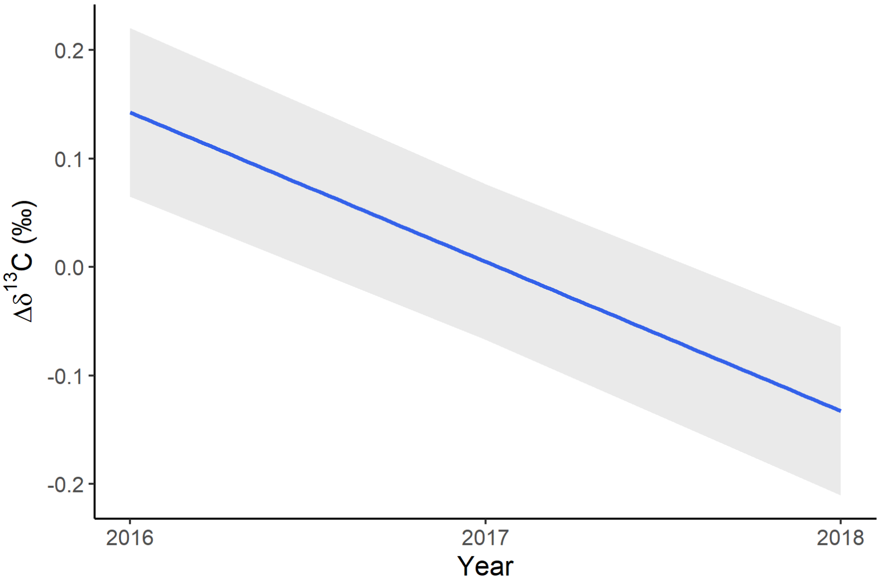


**Fig. S6** Effect of study year on $\delta^{13}C$ in wood of focal trees. The blue line is a linear mixed-effects model fit and the grey band shows a 95% confidence interval. See Table S6 for details on the fitted model.

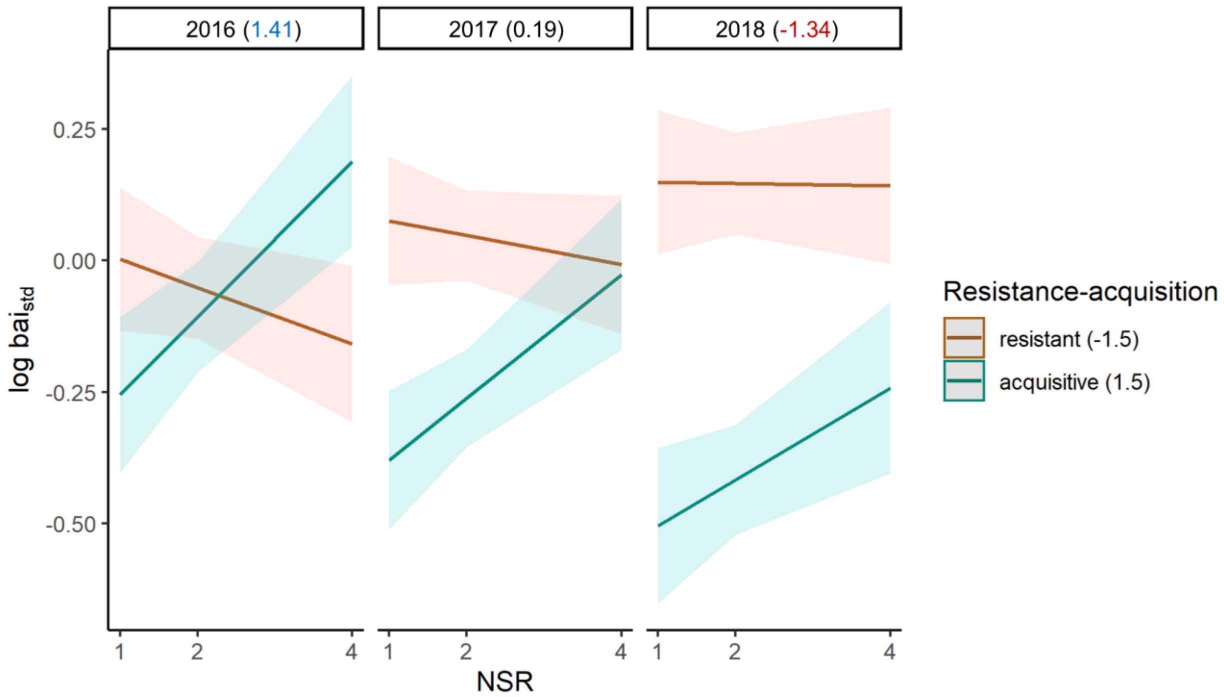


**Fig. S7** Modulation of the relationship between neighbourhood species richness (NSR), climate and growth by resistance-acquisition traits. Lines represent linear mixed-effects model fits and coloured bands show a 95% confidence interval. The models depict marginally significant, interactive effects of NSR and study year (2016-2018 with wet-to-dry climate, SPEI values in brackets) on growth of focal trees predicted for cavitation resistant (PC1 value of -1.5) and for acquisitive focal trees (PC1 value of 1.5). See Fig. 1 for details on the study design and Table S10 for details on the fitted model.

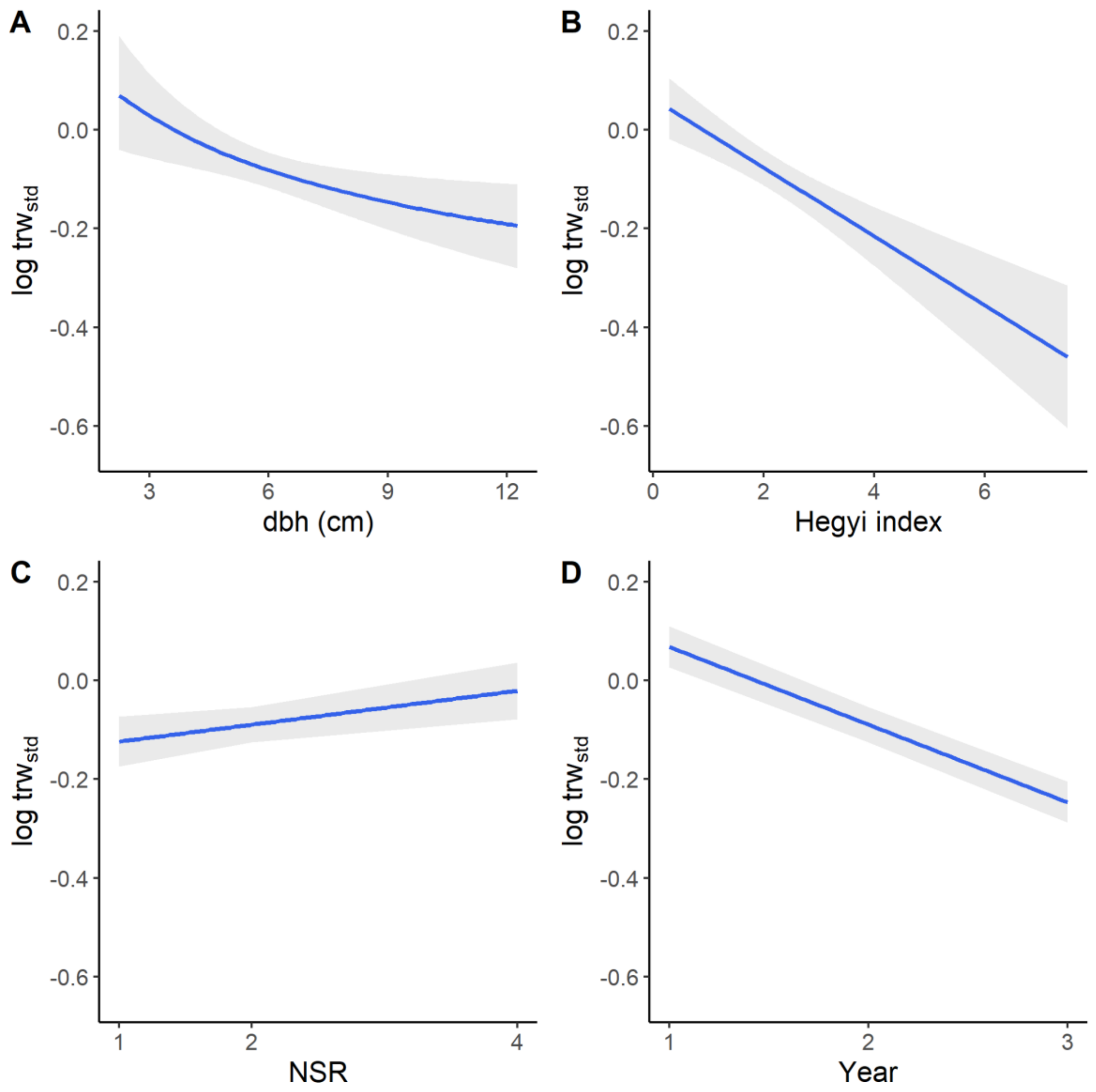


**Fig. S8** Effects of tree size (dbh), neighbourhood competition (Hegyi index), neighbourhood species richness (NSR) and study year on the logarithm of tree-ring width ($trw_{std}$) of focal trees. The blue lines are linear mixed-effects model fits and the grey bands show a 95% confidence interval.

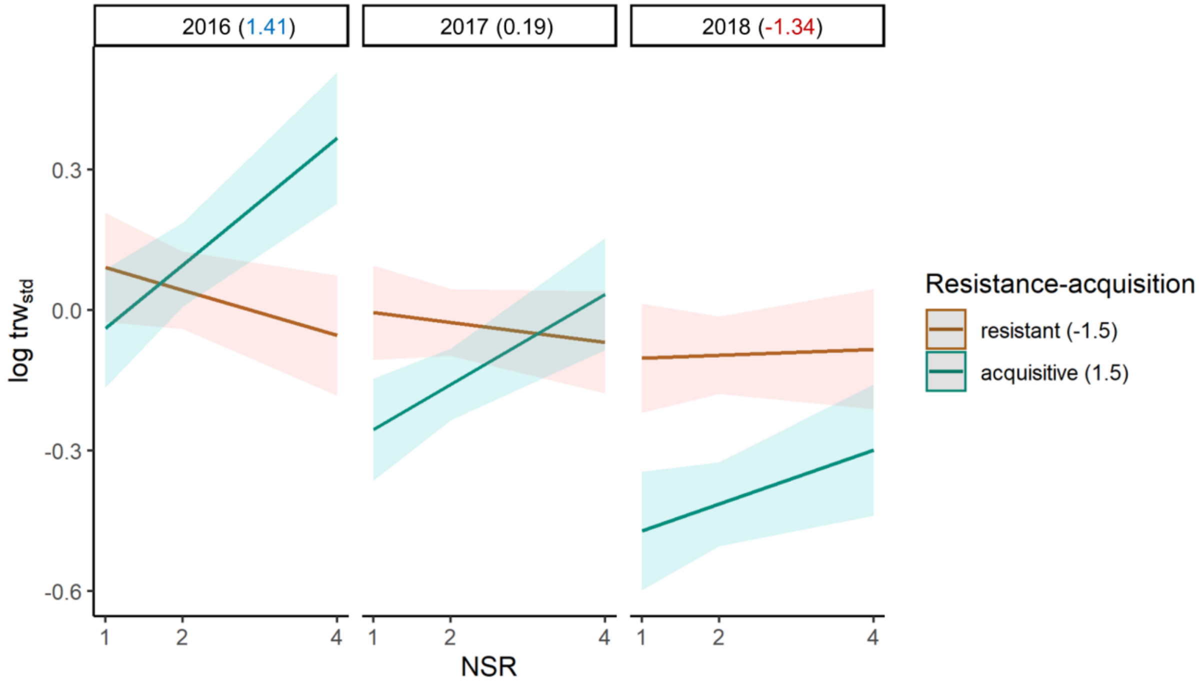


**Fig. S9** Modulation of the relationship between neighbourhood species richness (NSR), climate and growth by resistance-acquisition traits using tree-ring width ($trw_{std}$) of focal trees instead of basal area increment ($bai_{std}$) as indicator of growth. Lines represent linear mixed-effects model fits and coloured bands show a 95% confidence interval. The model depicts significant, interactive effects of NSR and study year (2016-2018 with wet-to-dry climate, SPEI values in brackets) on growth predicted for cavitation resistant (PC1 value of -1.5) and for acquisitive focal trees (PC1 value of 1.5) (NSR × year × focal tree resistance-acquisition traits, $t = -2.21$, $P = 0.027$). See Fig. 1 for details on the study design.

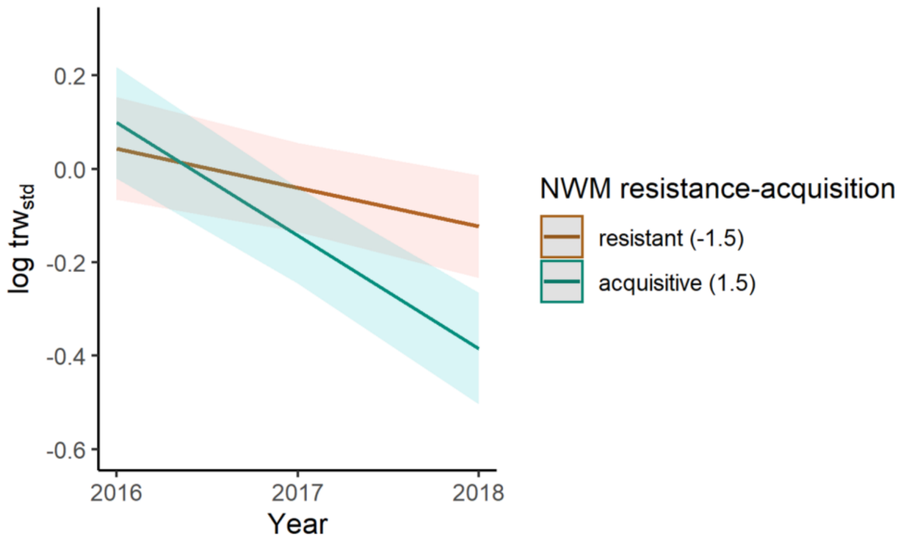


**Fig. S10** Modulation of the relationship between climate and growth by the neighbourhood-weighted mean (NWM) of resistance-acquisition traits using tree-ring width ($trw_{std}$) of focal trees instead of basal area increment ($bai_{std}$) as indicator of growth. Lines represent linear mixed-effects model fits and coloured bands show a 95% confidence interval. The model depicts a significant effect of study year (2016-2018 with wet-to-dry climate, SPEI values in brackets) on the logarithm of $trw_{std}$ predicted for a neighbourhood dominated by cavitation resistant (PC1 value of -1.5) and acquisitive species (PC1 value of 1.5) (year × NWM resistance-acquisition, $t = -2.90$, $P = 0.004$). See Fig. 1 for details on the study design.

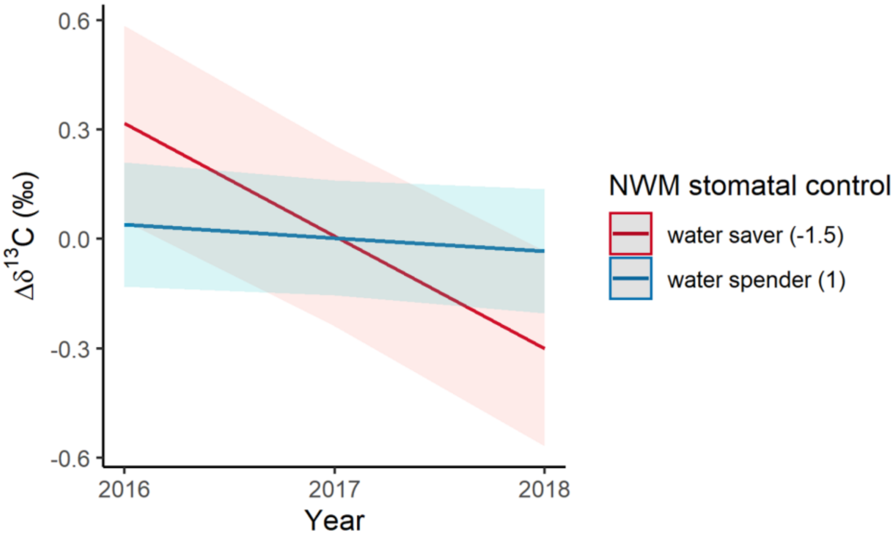


**Fig. S11** Modulation of the relationship between climate and $\delta^{13}C$ by the neighbourhood-weighted mean (NWM) of stomatal control traits. Lines represent linear mixed-effects model fits and coloured bands show a 95% confidence interval. The model depicts a significant effect of study year (2016-2018 with wet-to-dry climate, SPEI values in brackets) on $\delta^{13}C$ in wood of focal trees predicted for a water saver (PC2 value of -1.5) and for a water spender (PC2 value of 1.0) dominated neighbourhood. The sketch illustrates that the NWM of stomatal control traits (tree neighbourhood) modulates the relationship. See Fig. 1 for details on the study design and Table S16 for details on the fitted model.

**Table S1** The 40 broadleaved evergreen and deciduous tree species planted in BEF-China

| Species names | Family | Species code | Leaf habit | Site |
|---|---|---|---|---|
| *Acer davidii* | *Sapindaceae* | 27 | D | A |
| *Ailanthus altissima* | *Simaroubaceae* | 29 | D | B |
| ***Alniphyllum fortunei*** | ***Styracaceae*** | **30** | **D** | **B** |
| ***Betula luminifera*** | ***Betulaceae*** | **31** | **D** | **B** |
| ***Castanea henryi*** | ***Fagaceae*** | **1** | **D** | **A** |
| *Castanopsis carlesii* | *Fagaceae* | 10 | E | A |
| *Castanopsis eyrei* | *Fagaceae* | 13 | E | AB |
| ***Castanopsis fargesii*** | ***Fagaceae*** | **32** | **E** | **B** |
| *Castanopsis sclerophylla* | *Fagaceae* | 14 | E | AB |
| *Celtis biondii* | *Cannabaceae* | 33 | D | B |
| ***Choerospondias axillaris*** | ***Anacardiaceae*** | **4** | **D** | **A** |
| *Cinnamomum camphora* | *Lauraceae* | 17 | E | AB |
| *Cyclobalanopsis glauca* | *Fagaceae* | 11 | E | AB |
| *Cyclobalanopsis myrsinifolia* | *Fagaceae* | 9 | E | A |
| ***Daphniphyllum oldhamii*** | ***Daphniphyllaceae*** | **16** | **E** | **AB*** |
| *Diospyros japonica* | *Ebenaceae* | 15 | D | AB |
| ***Elaeocarpus chinensis*** | ***Elaeocarpaceae*** | **34** | **E** | **B** |
| *Elaeocarpus glabripetalus* | *Elaeocarpaceae* | 35 | E | B |
| ***Elaeocarpus japonicus*** | ***Elaeocarpaceae*** | **36** | **E** | **B** |
| ***Idesia polycarpa*** | ***Salicaceae*** | **37** | **D** | **B** |
| *Koelreuteria bipinnata* | *Sapindaceae* | 18 | D | A |
| ***Liquidambar formosana*** | ***Altingiaceae*** | **6** | **D** | **A** |
| ***Lithocarpus glaber*** | ***Fagaceae*** | **12** | **E** | **A*B** |
| *Machilus grijsii* | *Lauraceae* | 39 | E | B |
| *Machilus leptophylla* | *Lauraceae* | 41 | E | B |
| *Machilus thunbergii* | *Lauraceae* | 40 | E | B |
| *Manglietia fordiana* | *Magnoliaceae* | 42 | E | B |
| *Melia azedarach* | *Meliaceae* | 26 | D | A |
| *Meliosma flexuosa* | *Sabiaceae* | 38 | D | B |
| ***Nyssa sinensis*** | ***Cornaceae*** | **20** | **D** | **A** |
| *Phoebe bournei* | *Lauraceae* | 43 | E | B |
| ***Quercus acutissima*** | ***Fagaceae*** | **25** | **D** | **A** |
| *Quercus fabri* | *Fagaceae* | 24 | D | A |
| *Quercus phillyreoides* | *Fagaceae* | 44 | E | B |
| *Quercus serrata* | *Fagaceae* | 8 | D | A |
| *Rhus chinensis* | *Anacardiaceae* | 23 | D | A |
| *Sapindus saponaria* | *Sapindaceae* | 19 | D | A |
| ***Triadica cochinchinensis*** | ***Euphorbiaceae*** | **22** | **D** | **A** |
| *Triadica sebifera* | *Euphorbiaceae* | 21 | D | A |
| ***Schima superba*** | ***Theaceae*** | **3** | **E** | **A*B*** |

Note: Species from which tree cores were extracted are highlighted in bold (see Fig. 1 for the species selection). Shown are species and family names, the species identity codes used in Fig. 1, leaf habit (E, evergreen; D, Deciduous) and the site at which the species were planted. In case of species planted at both sites, asterisks indicate at which site the species was sampled. For more details on the tree species taxonomy, their characteristics and the experimental design see Bruelheide *et al.* (2014) and Huang *et al.* (2018).

**Table S2** Resistance-acquisition and stomatal control traits were used in this study (adapted from Schnabel *et al.* (2021)).

| Abbreviation | Trait description | Unit |
|---|---|---|
| Resistance | Water potential at which 50% initial conductivity is lost due to cavitation ($\Psi_{50}$) | MPa |
| SLA | Specific leaf area | $m^2$ $kg^{-1}$ |
| Leaf tough. | Leaf toughness | N $mm^{-1}$ |
| CN | Carbon to nitrogen ratio | Ratio |
| Stom. cond. | Modelled maximum stomatal conductance (CONMAXFIT) | Non-dimensional |
| Stom. density | Stomatal density | 1 $mm^{-2}$ |
| Stom. index | Product of Stom. density and stomatal size in $\mu m^2$ | ratio |
| Stom. reg. I | Stomatal regulation assessed as vapor pressure deficit (VPD) at modelled maximum stomatal conductance (VPDMAXFIT) | hPA |
| Stom. reg. II | Stomatal regulation assessed as VPD at the point of inflection of modelled stomatal conductance (VPDPOI) | hPA |

Note: Traits were measured in the BEF-China experiment and were used to calculate species level mean trait values by Kröber & Bruelheide (2014) and Kröber *et al.* (2014). See these studies and Schnabel *et al.* (2021) for detailed information on the individual traits and the two orthogonal drought-tolerance trait gradients they represent. Stomatal sensitivity (Stomatal regulation I and II) is inferred here from modelled $g_s$~VPD curves through extracting the point at which a species starts to lower its stomatal conductance (the VPD at maximum stomatal conductance, VPDMAXFIT) and the point where the slope of the curve turns from positive to negative (VPDPOI), which is a measure of how fast stomatal close under increasing VPD.

**Table S3** Description of competition indices.

| Index | Description |
|---|---|
| nhigher | Number of neighbours higher than the focal tree |
| relbah | Basal area of neighbours higher than the focal tree |
| relbab | Basal area of neighbours with higher basal area than the focal tree |
| hegyi | Hegyi index including all neighbours |
| hegyih | Hegyi index of neighbours higher than the focal tree |
| hegyib | Hegyi index of neighbours with higher basal area than the focal tree |
| hcom | Summed height of neighbours relative to the focal tree |

Notes: We modelled distance depended competition effects of neighbouring trees on focal trees using the Hegyi index (e.g. Mailly *et al.*, 2003) with the following formula when including all neighbours: $hegyi = \sum_{c=1}^{n} \frac{ba_c}{ba_t} * \frac{1}{d_{tc}}$; where *ba* is the basal area of either the focal tree *t* or its competitor *c* and $d_{tc}$ the distance between focal tree and neighbour. We subsequently adjusted this formula to only include those neighbours higher than the focal tree or those with a higher basal area than the focal tree. Tree basal area ($cm^2$, based on dbh), and in the case of multi-stemmed trees, the sum of basal areas of individual stems was used for calculating the respective indices.

**Table S4** Comparison of competition indices (Table S3) for the growth linear mixed-effects model (Table S4) against a null model without a competition index.

| Model | npar | AIC | BIC | logLik | deviance | Chisq | Df | Pr(>Chisq) |
|---|---|---|---|---|---|---|---|---|
| null | 6 | 912.56 | 942.05 | -450.28 | 900.56 | NA | NA | NA |
| nhigher | 7 | 914.23 | 948.64 | -450.11 | 900.23 | 0.33 | 1 | 0.57 |
| relbah | 7 | 913.18 | 947.59 | -449.59 | 899.18 | 1.05 | 0 | NA |
| relbab | 7 | 904.85 | 939.26 | -445.43 | 890.85 | 8.33 | 0 | NA |
| hegyi | 7 | 880.98 | 915.39 | -433.49 | 866.98 | 23.87 | 0 | NA |
| hegyih | 7 | 909.19 | 943.6 | -447.59 | 895.19 | 0 | 0 | NA |
| hegyib | 7 | 890.65 | 925.06 | -438.32 | 876.65 | 18.54 | 0 | NA |
| hcom | 7 | 914.14 | 948.55 | -450.07 | 900.14 | 0 | 0 | NA |

**Table S5** Best-fitting trait-independent linear mixed-effects model after model selection.

| | log($bai_{std}$) | | | | |
|---|---|---|---|---|---|
| *Predictors* | *Estimates* | *CI* | *Statistic* | *p* | *df* |
| (Intercept) | -0.19 | -0.29 – -0.10 | -4.18 | **<0.001** | 119.74 |
| dbh | 0.10 | 0.06 – 0.14 | 5.01 | **<0.001** | 273.26 |
| hegyi | -0.12 | -0.15 – -0.08 | -5.91 | **<0.001** | 328.58 |
| NSR | 0.04 | 0.01 – 0.08 | 2.29 | **0.024** | 120.00 |
| **Random Effects** | | | | | |
| $\sigma^2$ | 0.09 | | | | |
| $\tau_{00\ tag:(plot\_no:site)}$ | 0.07 | | | | |
| $\tau_{00\ plot\_no:site}$ | 0.02 | | | | |
| ICC | 0.51 | | | | |
| $N_{tag}$ | 336 | | | | |
| $N_{plot\_no}$ | 114 | | | | |
| $N_{site}$ | 2 | | | | |
| Observations | 1008 | | | | |
| Marginal $R^2$ / Conditional $R^2$ | 0.155 / 0.585 | | | | |

Note: Significant fixed effects printed in bold. Linear mixed-effects models (LMMs) fit with the packages lme4 (Bates *et al.*, 2015) and lmerTest (Kuznetsova *et al.*, 2017) in R using restricted maximum likelihood estimation (REML) and an α of 0.05 for reporting significant effects. Model tables including fixed and random effects as well as $R^2$ values were created using the sjPlot package (see Lüdecke (2021) for details). The model statistics, p-values, standard errors and confidence intervals (CI; 95%) were computed using Satterthwaite's approximation for degrees of freedom. 'Plot no' is the plot identifier and 'tag' the tree identifier. All analyses were conducted in R version 4.1.2 (R Core Team, 2021).

**Table S6** Best-fitting trait-independent linear mixed-effects model after model selection.

| | $\Delta\delta^{13}C$ | | | | |
|---|---|---|---|---|---|
| *Predictors* | *Estimates* | *CI* | *Statistic* | *p* | *df* |
| (Intercept) | 0.28 | 0.19 – 0.37 | 5.89 | **<0.001** | 256.95 |
| year int | -0.14 | -0.17 – -0.11 | -9.06 | **<0.001** | 671.00 |
| **Random Effects** | | | | | |
| $\sigma^2$ | 0.16 | | | | |
| $\tau_{00\ tag:(plot\_no:site)}$ | 0.29 | | | | |
| $\tau_{00\ plot\_no:site}$ | 0.03 | | | | |
| ICC | 0.67 | | | | |
| N $_{tag}$ | 336 | | | | |
| N $_{plot\_no}$ | 114 | | | | |
| N $_{site}$ | 2 | | | | |
| Observations | 1008 | | | | |
| Marginal $R^2$ / Conditional $R^2$ | 0.026 / 0.683 | | | | |

Note: Significant fixed effects printed in bold. Linear mixed-effects models (LMMs) fit with the packages lme4 (Bates *et al.*, 2015) and lmerTest (Kuznetsova *et al.*, 2017) in R using restricted maximum likelihood estimation (REML) and an $\alpha$ of 0.05 for reporting significant effects. Model tables including fixed and random effects as well as $R^2$ values were created using the sjPlot package (see Lüdecke (2021) for details). The model statistics, p-values, standard errors and confidence intervals (CI; 95%) were computed using Satterthwaite's approximation for degrees of freedom. 'Plot no' is the plot identifier and 'tag' the tree identifier. All analyses were conducted in R version 4.1.2 (R Core Team, 2021).

**Table S7** Comparison of competition indices (Table S3) for the $\delta^{13}C$ linear mixed-effects model (Table S6) against a null model without a competition index.

| Models | npar | AIC | BIC | logLik | deviance | Chisq | Df | Pr(>Chisq) |
|---|---|---|---|---|---|---|---|---|
| null | 5 | 1650.47 | 1675.04 | -820.23 | 1640.47 | NA | NA | NA |
| nhigher | 6 | 1652.22 | 1681.71 | -820.11 | 1640.22 | 0.25 | 1 | 0.62 |
| relbah | 6 | 1650.80 | 1680.30 | -819.40 | 1638.80 | 1.41 | 0 | NA |
| relbab | 6 | 1652.09 | 1681.59 | -820.05 | 1640.09 | 0.00 | 0 | NA |
| hegyi | 6 | 1651.34 | 1680.83 | -819.67 | 1639.34 | 0.75 | 0 | NA |
| hegyih | 6 | 1650.72 | 1680.22 | -819.36 | 1638.72 | 0.61 | 0 | NA |
| hegyib | 6 | 1652.38 | 1681.88 | -820.19 | 1640.38 | 0.00 | 0 | NA |
| hcom | 6 | 1652.44 | 1681.94 | -820.22 | 1640.44 | 0.00 | 0 | NA |

**Table S8** Best-fitting linear mixed-effects model of focal tree resistance-acquisition traits after model selection.

| | **log(bai$_{std}$)** | | | | |
|---|---|---|---|---|---|
| *Predictors* | *Estimates* | *CI* | *Statistic* | *p* | *df* |
| (Intercept) | -0.15 | -0.25 – -0.06 | -3.13 | **0.002** | 178.59 |
| dbh | 0.12 | 0.08 – 0.16 | 5.80 | **<0.001** | 276.94 |
| hegyi | -0.12 | -0.16 – -0.08 | -6.06 | **<0.001** | 324.39 |
| NSR | 0.04 | 0.01 – 0.07 | 2.40 | **0.018** | 116.73 |
| resistance-acquisition | -0.02 | -0.11 – 0.08 | -0.31 | 0.758 | 193.94 |
| year int | -0.02 | -0.04 – 0.00 | -1.79 | 0.074 | 670.00 |
| NSR * resistance-acquisition | 0.04 | 0.01 – 0.07 | 2.45 | **0.015** | 160.99 |
| resistance-acquisition * year int | -0.08 | -0.10 – -0.05 | -6.84 | **<0.001** | 670.00 |
| **Random Effects** | | | | | |
| $\sigma^2$ | 0.08 | | | | |
| $\tau_{00\ tag:(plot\_no:site)}$ | 0.07 | | | | |
| $\tau_{00\ plot\_no:site}$ | 0.01 | | | | |
| ICC | 0.52 | | | | |
| N $_{tag}$ | 336 | | | | |
| N $_{plot\_no}$ | 114 | | | | |
| N $_{site}$ | 2 | | | | |
| Observations | 1008 | | | | |
| Marginal $R^2$ / Conditional $R^2$ | 0.200 / 0.612 | | | | |

Note: Significant fixed effects printed in bold. Linear mixed-effects models (LMMs) fit with the packages lme4 (Bates *et al.*, 2015) and lmerTest (Kuznetsova *et al.*, 2017) in R using restricted maximum likelihood estimation (REML) and an α of 0.05 for reporting significant effects. Model tables including fixed and random effects as well as $R^2$ values were created using the sjPlot package (see Lüdecke (2021) for details). The model statistics, p-values, standard errors and confidence intervals (CI; 95%) were computed using Satterthwaite's approximation for degrees of freedom. 'Plot no' is the plot identifier and 'tag' the tree identifier. All analyses were conducted in R version 4.1.2 (R Core Team, 2021).

**Table S9** Best-fitting linear mixed-effects model of neighbour resistance-acquisition traits after model selection.

| | log($bai_{std}$) | | | | |
|---|---|---|---|---|---|
| *Predictors* | *Estimates* | *CI* | *Statistic* | *p* | *df* |
| (Intercept) | -0.15 | -0.25 – -0.05 | -2.88 | **0.004** | 176.14 |
| dbh | 0.12 | 0.08 – 0.16 | 5.58 | **<0.001** | 282.81 |
| hegyi | -0.12 | -0.16 – -0.08 | -6.07 | **<0.001** | 324.78 |
| NSR | 0.04 | 0.00 – 0.07 | 2.12 | **0.036** | 115.21 |
| year int | -0.02 | -0.04 – 0.00 | -1.75 | 0.080 | 670.00 |
| resistance-acquisition | 0.03 | -0.03 – 0.10 | 1.08 | 0.282 | 490.68 |
| year int * resistance-acquisition | -0.05 | -0.07 – -0.02 | -4.17 | **<0.001** | 670.00 |
| **Random Effects** | | | | | |
| $\sigma^2$ | 0.09 | | | | |
| $\tau_{00\ tag:(plot\_no:site)}$ | 0.07 | | | | |
| $\tau_{00\ plot\_no:site}$ | 0.02 | | | | |
| ICC | 0.51 | | | | |
| N tag | 336 | | | | |
| N plot_no | 114 | | | | |
| N site | 2 | | | | |
| Observations | 1008 | | | | |
| Marginal $R^2$ / Conditional $R^2$ | 0.173 / 0.595 | | | | |

Note: Significant fixed effects printed in bold. Linear mixed-effects models (LMMs) fit with the packages lme4 (Bates *et al.*, 2015) and lmerTest (Kuznetsova *et al.*, 2017) in R using restricted maximum likelihood estimation (REML) and an α of 0.05 for reporting significant effects. Model tables including fixed and random effects as well as $R^2$ values were created using the sjPlot package (see Lüdecke (2021) for details). The model statistics, p-values, standard errors and confidence intervals (CI; 95%) were computed using Satterthwaite's approximation for degrees of freedom. 'Plot no' is the plot identifier and 'tag' the tree identifier. All analyses were conducted in R version 4.1.2 (R Core Team, 2021).

**Table S10** Best-fitting linear mixed-effects model of focal tree stomatal control traits after model selection.

| | **log($bai_{std}$)** | | | | |
|---|---|---|---|---|---|
| *Predictors* | *Estimates* | *CI* | *Statistic* | *p* | *df* |
| (Intercept) | -0.16 | -0.26 – -0.05 | -3.00 | **0.003** | 177.35 |
| dbh | 0.10 | 0.06 – 0.14 | 5.02 | **<0.001** | 274.02 |
| hegyi | -0.12 | -0.15 – -0.08 | -5.89 | **<0.001** | 327.59 |
| NSR | 0.04 | 0.01 – 0.08 | 2.29 | **0.024** | 118.62 |
| year int | -0.02 | -0.04 – 0.00 | -1.77 | 0.078 | 670.00 |
| stomatal control | 0.10 | 0.04 – 0.17 | 3.37 | **0.001** | 503.88 |
| year int * stomatal control | -0.06 | -0.08 – -0.04 | -5.10 | **<0.001** | 670.00 |
| **Random Effects** | | | | | |
| $\sigma^2$ | 0.08 | | | | |
| $\tau_{00\ tag:(plot\_no:site)}$ | 0.07 | | | | |
| $\tau_{00\ plot\_no:site}$ | 0.02 | | | | |
| ICC | 0.52 | | | | |
| $N_{tag}$ | 336 | | | | |
| $N_{plot\_no}$ | 114 | | | | |
| $N_{site}$ | 2 | | | | |
| Observations | 1008 | | | | |
| Marginal $R^2$ / Conditional $R^2$ | 0.166 / 0.603 | | | | |

Note: Significant fixed effects printed in bold. Linear mixed-effects models (LMMs) fit with the packages lme4 (Bates *et al.*, 2015) and lmerTest (Kuznetsova *et al.*, 2017) in R using restricted maximum likelihood estimation (REML) and an $\alpha$ of 0.05 for reporting significant effects. Model tables including fixed and random effects as well as $R^2$ values were created using the sjPlot package (see Lüdecke (2021) for details). The model statistics, p-values, standard errors and confidence intervals (CI; 95%) were computed using Satterthwaite's approximation for degrees of freedom. 'Plot no' is the plot identifier and 'tag' the tree identifier. All analyses were conducted in R version 4.1.2 (R Core Team, 2021).

**Table S11** Best-fitting linear mixed-effects model of neighbour stomatal control traits after model selection.

| | $\log(bai_{std})$ | | | | |
|---|---|---|---|---|---|
| *Predictors* | *Estimates* | *CI* | *Statistic* | *p* | *df* |
| (Intercept) | -0.15 | -0.26 – -0.05 | -2.95 | **0.004** | 179.65 |
| dbh | 0.10 | 0.06 – 0.14 | 4.95 | **<0.001** | 275.70 |
| hegyi | -0.12 | -0.16 – -0.08 | -5.92 | **<0.001** | 328.73 |
| NSR | 0.04 | 0.00 – 0.07 | 2.23 | **0.027** | 119.44 |
| year int | -0.02 | -0.04 – 0.00 | -1.74 | 0.082 | 670.00 |
| stomatal control | 0.08 | 0.02 – 0.14 | 2.57 | **0.011** | 502.83 |
| year int * stomatal control | -0.03 | -0.06 – -0.01 | -3.04 | **0.002** | 670.00 |
| **Random Effects** | | | | | |
| $\sigma^2$ | 0.09 | | | | |
| $\tau_{00\ tag:(plot\_no:site)}$ | 0.07 | | | | |
| $\tau_{00\ plot\_no:site}$ | 0.02 | | | | |
| ICC | 0.52 | | | | |
| $N_{tag}$ | 336 | | | | |
| $N_{plot\_no}$ | 114 | | | | |
| $N_{site}$ | 2 | | | | |
| Observations | 1008 | | | | |
| Marginal $R^2$ / Conditional $R^2$ | 0.160 / 0.593 | | | | |

Note: Significant fixed effects printed in bold. Linear mixed-effects models (LMMs) fit with the packages lme4 (Bates *et al.*, 2015) and lmerTest (Kuznetsova *et al.*, 2017) in R using restricted maximum likelihood estimation (REML) and an α of 0.05 for reporting significant effects. Model tables including fixed and random effects as well as $R^2$ values were created using the sjPlot package (see Lüdecke (2021) for details). The model statistics, p-values, standard errors and confidence intervals (CI; 95%) were computed using Satterthwaite's approximation for degrees of freedom. 'Plot no' is the plot identifier and 'tag' the tree identifier. All analyses were conducted in R version 4.1.2 (R Core Team, 2021).

**Table S12** Best-fitting linear mixed-effects model of focal tree resistance-acquisition traits after model selection.

| | $\Delta\delta^{13}C$ | | | | |
|---|---|---|---|---|---|
| *Predictors* | *Estimates* | *CI* | *Statistic* | *p* | *df* |
| (Intercept) | 0.16 | 0.08 – 0.24 | 3.91 | **<0.001** | 140.98 |
| year cat [2017] | -0.19 | -0.25 – -0.13 | -6.21 | **<0.001** | 668.00 |
| year cat [2018] | -0.28 | -0.33 – -0.22 | -9.14 | **<0.001** | 668.00 |
| resistance-acquisition | -0.06 | -0.14 – 0.02 | -1.51 | 0.133 | 192.66 |
| year cat [2017] * resistance-acquisition | 0.11 | 0.05 – 0.16 | 3.51 | **<0.001** | 668.00 |
| year cat [2018] * resistance-acquisition | 0.06 | -0.00 – 0.11 | 1.84 | 0.066 | 668.00 |
| **Random Effects** | | | | | |
| $\sigma^2$ | 0.15 | | | | |
| $\tau_{00\ tag:(plot\_no:site)}$ | 0.29 | | | | |
| $\tau_{00\ plot\_no:site}$ | 0.04 | | | | |
| ICC | 0.68 | | | | |
| $N_{tag}$ | 336 | | | | |
| $N_{plot\_no}$ | 114 | | | | |
| $N_{site}$ | 2 | | | | |
| Observations | 1008 | | | | |
| Marginal $R^2$ / Conditional $R^2$ | 0.031 / 0.690 | | | | |

Note: Significant fixed effects printed in bold. Linear mixed-effects models (LMMs) fit with the packages lme4 (Bates *et al.*, 2015) and lmerTest (Kuznetsova *et al.*, 2017) in R using restricted maximum likelihood estimation (REML) and an α of 0.05 for reporting significant effects. Model tables including fixed and random effects as well as $R^2$ values were created using the sjPlot package (see Lüdecke (2021) for details). The model statistics, p-values, standard errors and confidence intervals (CI; 95%) were computed using Satterthwaite's approximation for degrees of freedom. 'Plot no' is the plot identifier and 'tag' the tree identifier. All analyses were conducted in R version 4.1.2 (R Core Team, 2021).

**Table S13** Best-fitting linear mixed-effects model of neighbour resistance-acquisition traits after model selection.

| | | | $\Delta\delta^{13}C$ | | |
|---|---|---|---|---|---|
| *Predictors* | *Estimates* | *CI* | *Statistic* | *p* | *df* |
| (Intercept) | 0.16 | 0.08 – 0.24 | 3.92 | **<0.001** | 139.09 |
| year cat [2017] | -0.19 | -0.25 – -0.13 | -6.24 | **<0.001** | 668.00 |
| year cat [2018] | -0.28 | -0.33 – -0.22 | -9.19 | **<0.001** | 668.00 |
| resistance-acquisition | -0.05 | -0.13 – 0.03 | -1.22 | 0.224 | 151.57 |
| year cat [2017] * resistance-acquisition | 0.13 | 0.07 – 0.19 | 4.35 | **<0.001** | 668.00 |
| year cat [2018] * resistance-acquisition | 0.07 | 0.01 – 0.13 | 2.27 | **0.023** | 668.00 |
| **Random Effects** | | | | | |
| $\sigma^2$ | 0.15 | | | | |
| $\tau_{00\ tag:(plot\_no:site)}$ | 0.29 | | | | |
| $\tau_{00\ plot\_no:site}$ | 0.03 | | | | |
| ICC | 0.68 | | | | |
| $N_{tag}$ | 336 | | | | |
| $N_{plot\_no}$ | 114 | | | | |
| $N_{site}$ | 2 | | | | |
| Observations | 1008 | | | | |
| Marginal $R^2$ / Conditional $R^2$ | 0.033 / 0.692 | | | | |

Note: Significant fixed effects printed in bold. Linear mixed-effects models (LMMs) fit with the packages lme4 (Bates *et al.*, 2015) and lmerTest (Kuznetsova *et al.*, 2017) in R using restricted maximum likelihood estimation (REML) and an α of 0.05 for reporting significant effects. Model tables including fixed and random effects as well as $R^2$ values were created using the sjPlot package (see Lüdecke (2021) for details). The model statistics, p-values, standard errors and confidence intervals (CI; 95%) were computed using Satterthwaite's approximation for degrees of freedom. 'Plot no' is the plot identifier and 'tag' the tree identifier. All analyses were conducted in R version 4.1.2 (R Core Team, 2021).

**Table S14** Best-fitting linear mixed-effects model of focal tree stomatal control traits after model selection.

| | $\Delta\delta^{13}C$ | | | | |
|---|---|---|---|---|---|
| *Predictors* | *Estimates* | *CI* | *Statistic* | *p* | *df* |
| (Intercept) | 0.36 | 0.16 – 0.56 | 3.61 | **<0.001** | 235.28 |
| NSR | -0.04 | -0.11 – 0.04 | -0.93 | 0.351 | 239.44 |
| year int | -0.15 | -0.21 – -0.09 | -4.76 | **<0.001** | 668.00 |
| stomatal control | -0.29 | -0.48 – -0.09 | -2.84 | **0.005** | 228.98 |
| NSR * year int | 0.01 | -0.02 – 0.03 | 0.51 | 0.610 | 668.00 |
| NSR * stomatal control | 0.09 | 0.01 – 0.16 | 2.26 | **0.024** | 315.29 |
| year int * stomatal control | 0.12 | 0.06 – 0.18 | 3.73 | **<0.001** | 668.00 |
| (NSR * year int) * stomatal control | -0.03 | -0.06 – -0.01 | -2.66 | **0.008** | 668.00 |
| **Random Effects** | | | | | |
| $\sigma^2$ | 0.15 | | | | |
| $\tau_{00\ tag:(plot\_no:site)}$ | 0.29 | | | | |
| $\tau_{00\ plot\_no:site}$ | 0.03 | | | | |
| ICC | 0.68 | | | | |
| $N_{tag}$ | 336 | | | | |
| $N_{plot\_no}$ | 114 | | | | |
| $N_{site}$ | 2 | | | | |
| Observations | 1008 | | | | |
| Marginal $R^2$ / Conditional $R^2$ | 0.034 / 0.691 | | | | |

Note: Significant fixed effects printed in bold. Linear mixed-effects models (LMMs) fit with the packages lme4 (Bates *et al.*, 2015) and lmerTest (Kuznetsova *et al.*, 2017) in R using restricted maximum likelihood estimation (REML) and an α of 0.05 for reporting significant effects. Model tables including fixed and random effects as well as $R^2$ values were created using the sjPlot package (see Lüdecke (2021) for details). The model statistics, p-values, standard errors and confidence intervals (CI; 95%) were computed using Satterthwaite's approximation for degrees of freedom. 'Plot no' is the plot identifier and 'tag' the tree identifier. All analyses were conducted in R version 4.1.2 (R Core Team, 2021).

**Table S15** Best-fitting linear mixed-effects model of neighbour stomatal control traits after model selection.

| | $\Delta\delta^{13}C$ | | | | |
|---|---|---|---|---|---|
| *Predictors* | *Estimates* | *CI* | *Statistic* | *p* | *df* |
| (Intercept) | 0.28 | 0.19 – 0.37 | 5.90 | **<0.001** | 250.05 |
| year int | -0.14 | -0.17 – -0.11 | -9.13 | **<0.001** | 670.00 |
| stomatal control | -0.10 | -0.20 – -0.01 | -2.21 | **0.028** | 291.89 |
| year int * stomatal control | 0.05 | 0.02 – 0.08 | 3.43 | **0.001** | 670.00 |
| **Random Effects** | | | | | |
| $\sigma^2$ | 0.15 | | | | |
| $\tau_{00}$ tag:(plot_no:site) | 0.29 | | | | |
| $\tau_{00}$ plot_no:site | 0.04 | | | | |
| ICC | 0.68 | | | | |
| N tag | 336 | | | | |
| N plot_no | 114 | | | | |
| N site | 2 | | | | |
| Observations | 1008 | | | | |
| Marginal $R^2$ / Conditional $R^2$ | 0.029 / 0.689 | | | | |

Note: Significant fixed effects printed in bold. Linear mixed-effects models (LMMs) fit with the packages lme4 (Bates *et al.*, 2015) and lmerTest (Kuznetsova *et al.*, 2017) in R using restricted maximum likelihood estimation (REML) and an α of 0.05 for reporting significant effects. Model tables including fixed and random effects as well as $R^2$ values were created using the sjPlot package (see Lüdecke (2021) for details). The model statistics, p-values, standard errors and confidence intervals (CI; 95%) were computed using Satterthwaite's approximation for degrees of freedom. 'Plot no' is the plot identifier and 'tag' the tree identifier. All analyses were conducted in R version 4.1.2 (R Core Team, 2021).

**Table S16** Linear mixed-effects model of focal tree resistance-acquisition traits that still includes the marginally significant 3-way interaction between year, neighbourhood species richness (NSR) and resistance-acquisition traits.

| | log($bai_{std}$) | | | | |
|---|---|---|---|---|---|
| *Predictors* | *Estimates* | *CI* | *Statistic* | *p* | *df* |
| (Intercept) | -0.16 | -0.28 – -0.03 | -2.43 | **0.015** | 425.58 |
| dbh | 0.12 | 0.08 – 0.16 | 5.80 | **<0.001** | 276.94 |
| hegyi | -0.12 | -0.16 – -0.08 | -6.06 | **<0.001** | 324.39 |
| NSR | 0.04 | -0.01 – 0.09 | 1.68 | 0.093 | 428.37 |
| year int | -0.02 | -0.06 – 0.03 | -0.80 | 0.422 | 668.00 |
| resistance-acquisition | -0.09 | -0.21 – 0.04 | -1.35 | 0.177 | 451.55 |
| NSR * year int | -0.00 | -0.02 – 0.02 | -0.05 | 0.963 | 668.00 |
| NSR * resistance-acquisition | 0.07 | 0.02 – 0.12 | 2.95 | **0.003** | 566.08 |
| year int * resistance-acquisition | -0.04 | -0.09 – 0.01 | -1.68 | 0.093 | 668.00 |
| (NSR * year int) * resistance-acquisition | -0.02 | -0.03 – 0.00 | -1.75 | 0.080 | 668.00 |
| **Random Effects** | | | | | |
| $\sigma^2$ | 0.08 | | | | |
| $\tau_{00\ tag:(plot\_no:site)}$ | 0.07 | | | | |
| $\tau_{00\ plot\_no:site}$ | 0.01 | | | | |
| ICC | 0.52 | | | | |
| N tag | 336 | | | | |
| N plot_no | 114 | | | | |
| N site | 2 | | | | |
| Observations | 1008 | | | | |
| Marginal $R^2$ / Conditional $R^2$ | 0.201 / 0.613 | | | | |

Note: Significant fixed effects printed in bold. Linear mixed-effects models (LMMs) fit with the packages lme4 (Bates *et al.*, 2015) and lmerTest (Kuznetsova *et al.*, 2017) in R using restricted maximum likelihood estimation (REML) and an α of 0.05 for reporting significant effects. Model tables including fixed and random effects as well as $R^2$ values were created using the sjPlot package (see Lüdecke (2021) for details). The model statistics, p-values, standard errors and confidence intervals (CI; 95%) were computed using Satterthwaite's approximation for degrees of freedom. ‘Plot no’ is the plot identifier and ‘tag’ the tree identifier. All analyses were conducted in R version 4.1.2 (R Core Team, 2021).

### Supplementary Method 1 – focal tree selection

Focal trees were selected outside the plot's inventoried core area to avoid interference with other projects and not near plot borders unless the adjacent plot contained species for which complete trait information existed. We defined NSR as the realised neighbourhood species richness at sampling (2019). In a few cases (*Idesia polycarpa*, *Triadica cochinchinensis*), we used the original plot-level richness at the time of planting due to the high mortality of focal and neighbour species. For NSR=1 and NSR=2, we selected trees only from plots with the respective plot-level design richness (monocultures and 2-species mixtures). The random planting design of BEF-China substantially reduced the likelihood of finding high-diversity neighbourhoods. At NSR=4, we thus also selected plots with a higher plot-level design richness, i.e. 8- and 16-species mixtures in this order of preference. Sampling neighbourhoods with four species only in 4-species mixtures was impossible due to the high mortality of neighbour species. We used a focal tree threshold diameter at breast height (dbh, at 1.3m) of > 8cm (in a few cases ~6cm) to avoid damaging the trees when extracting cores for analysis.

In contrast to a former study which did not control for tree size and canopy position during sampling and consequently found strong shading effects on $\delta^{13}C$ (Jansen *et al.*, 2021), we aimed at maximizing the climate-induced water availability signal in $\delta^{13}C$ through focusing on (co-dominant) trees with little shading by their neighbourhood (see also Grossiord *et al.*, 2014; Jucker *et al.*, 2017). We did this in two ways. First, our selection of species with mid- to fast growth rates and the use of a minimum dbh (see above) resulted in a sample of (co-)dominant individuals. Second, to further reduce shading effects on focal trees and to simultaneously ensure the same sample size across species and NSR levels, we carried out an *a priori* selection of those focal trees that experienced least shading by their neighbourhood. For that purpose, those 7 focal trees per species, NSR-level and site with the highest sum of relative heights of neighbours (height of focal tree minus height of neighbour) (see above) were selected for statistical analysis. We constrained this selection to keep a minimum of two trees per plot.

### Supplementary Method 2 – trait-independent neighbourhood models

We included focal tree size as the log-transformed initial dbh (at the end of 2015) before our analysis period (2016-2018), which we estimated using tree-ring width and the trees' dbh recorded in 2019. We then tested eight neighbourhood competition indices (Table S3) and selected the best-performing one via the Akaike Information Criterion (AIC). We accounted for differences in average annual climatic conditions using year (2016-wet, 2017-intermediate and 2018-dry, Fig. 1A) as a fixed effect, coded as an integer variable as we expected linear trends. We also checked for non-linear behaviour through alternative models with year as a categorical fixed effect; in the latter case we report F values (based on type-III sum of squares) instead of t values. We used years rather than SPEI-values as we only examined three years with a clear gradient in climate-induced water availability. Reported relationships would be the same if using SPEI values (checked via alternative LMMs with SPEI-values as a fixed effect). We selected the most parsimonious random effect structure using the step function in lmerTest ($\alpha$=0.2, a conservative choice), which retained tree identity and plot but not site as nested random effects for models to explain growth and $\delta^{13}C$. Reported relationships were insensitive to the inclusion or exclusion of site as random effect as it did not explain significant variation in any of the examined models. That observed relationships did not differ between sites is also in line with the results of our more detailed analysis in this regard using the species *Schima superba* (see Supplementary Analysis 1). We kept this random effect structure in all further analyses. Finally, we selected the most parsimonious trait-independent neighbourhood model through backward elimination of fixed effects ($\alpha$=0.05).

### Supplementary Analysis 1 – potential site-specific differences

To test for potential differences between site A and B of the BEF-China experiment (Bruelheide *et al.*, 2014), we conducted a separate analysis of *Schima superba*, the species sampled at both sites (Fig. 1, Table S1). We used the same linear mixed-effects model (LMMs) structure as in the main analyses but included site as fixed and not as random effect. LMMs thus modelled growth and $\delta^{13}C$ in response to focal tree size, competition by neighbours, climate, neighbourhood species richness (NSR) and site. We also included the 3-way interaction between climate × NSR × site as well as all potential 2-way interactions as fixed effects. We did not include species traits as we only examined one species. These analyses indicated that growth and $\delta^{13}C$ responses did not differ between sites (see Tables S17,18 for the full models).

**Table S17** Full linear mixed-effects model for growth of the species *Schima superba* exploring the influence of experimental site (A or B).

| | log($bai_{std}$) | | | | |
|---|---|---|---|---|---|
| *Predictors* | *Estimates* | *CI* | *Statistic* | *p* | *df* |
| (Intercept) | 0.20 | -0.15 – 0.54 | 1.16 | 0.256 | 30.49 |
| dbh | 0.22 | 0.13 – 0.32 | 4.83 | **<0.001** | 31.38 |
| hegyi | -0.10 | -0.19 – -0.01 | -2.23 | **0.033** | 33.82 |
| NSR | -0.09 | -0.22 – 0.04 | -1.46 | 0.153 | 34.34 |
| year int | -0.15 | -0.24 – -0.05 | -3.15 | **0.002** | 80.00 |
| site [B] | -0.18 | -0.72 – 0.35 | -0.74 | 0.472 | 13.16 |
| NSR * year int | 0.05 | 0.01 – 0.08 | 2.80 | **0.006** | 80.00 |
| NSR * site [B] | 0.05 | -0.13 – 0.23 | 0.60 | 0.558 | 16.30 |
| year int * site [B] | 0.09 | -0.04 – 0.22 | 1.37 | 0.174 | 80.00 |
| (NSR * year int) * site [B] | -0.02 | -0.07 – 0.03 | -0.74 | 0.459 | 80.00 |
| **Random Effects** | | | | | |
| $\sigma^2$ | 0.02 | | | | |
| $\tau_{00\ tag:plot\_no}$ | 0.05 | | | | |
| $\tau_{00\ plot\_no}$ | 0.01 | | | | |
| ICC | 0.72 | | | | |
| $N_{tag}$ | 42 | | | | |
| $N_{plot\_no}$ | 16 | | | | |
| Observations | 126 | | | | |
| Marginal $R^2$ / Conditional $R^2$ | 0.468 / 0.854 | | | | |

Note: Significant fixed effects printed in bold. Linear mixed-effects models (LMMs) fit with the packages lme4 (Bates *et al.*, 2015) and lmerTest (Kuznetsova *et al.*, 2017) in R using restricted maximum likelihood estimation (REML) and an α of 0.05 for reporting significant effects. Model tables including fixed and random effects as well as $R^2$ values were created using the sjPlot package (see Lüdecke (2021) for details). The model statistics, p-values, standard errors and confidence intervals (CI; 95%) were computed using Satterthwaite's approximation for degrees of freedom. All analyses were conducted in R version 4.1.2 (R Core Team, 2021).

**Table S18** Full linear mixed-effects model for growth of the species *Schima superba* exploring the influence of experimental site (A or B).

| | **Δδ¹³C** | | | | |
|---|---|---|---|---|---|
| *Predictors* | *Estimates* | *CI* | *Statistic* | *p* | *df* |
| (Intercept) | 0.20 | -0.77 – 1.17 | 0.42 | 0.676 | 28.76 |
| dbh | 0.13 | -0.09 – 0.35 | 1.18 | 0.247 | 27.81 |
| hegyi | 0.11 | -0.11 – 0.33 | 1.04 | 0.305 | 30.79 |
| NSR | 0.03 | -0.33 – 0.39 | 0.16 | 0.871 | 32.03 |
| year int | -0.19 | -0.45 – 0.07 | -1.45 | 0.150 | 80.00 |
| site [B] | 0.66 | -0.90 – 2.22 | 0.90 | 0.384 | 16.14 |
| NSR * year int | 0.02 | -0.08 – 0.12 | 0.42 | 0.679 | 80.00 |
| NSR * site [B] | -0.28 | -0.82 – 0.25 | -1.11 | 0.283 | 18.62 |
| year int * site [B] | -0.05 | -0.42 – 0.32 | -0.26 | 0.795 | 80.00 |
| (NSR * year int) * site [B] | 0.04 | -0.10 – 0.18 | 0.64 | 0.525 | 80.00 |
| **Random Effects** | | | | | |
| $\sigma^2$ | 0.16 | | | | |
| $\tau_{00\ tag:plot\_no}$ | 0.21 | | | | |
| $\tau_{00\ plot\_no}$ | 0.15 | | | | |
| ICC | 0.69 | | | | |
| $N_{tag}$ | 42 | | | | |
| $N_{plot\_no}$ | 16 | | | | |
| Observations | 126 | | | | |
| Marginal $R^2$ / Conditional $R^2$ | 0.078 / 0.713 | | | | |

Note: Significant fixed effects printed in bold. Linear mixed-effects models (LMMs) fit with the packages lme4 (Bates *et al.*, 2015) and lmerTest (Kuznetsova *et al.*, 2017) in R using restricted maximum likelihood estimation (REML) and an α of 0.05 for reporting significant effects. Model tables including fixed and random effects as well as $R^2$ values were created using the sjPlot package (see Lüdecke (2021) for details). The model statistics, p-values, standard errors and confidence intervals (CI; 95%) were computed using Satterthwaite's approximation for degrees of freedom. All analyses were conducted in R version 4.1.2 (R Core Team, 2021).

### Supplementary Discussion 1 – coordination of drought-tolerance traits

We observed species responses in wet to dry years along the resistance-acquisition gradient consistent with the current understanding of a trade-off between high cavitation resistance (low $\Psi_{50}$) and acquisitive resource use in tropical tree species (Guillemot *et al.*, 2022). Similarly, Reich (2014) concluded that acquisitive species thrive under optimal conditions due to their high capacity to transport and store water, while resistant species have slower resource economics but are less vulnerable to drought. In line with this notion, also the specific hydraulic conductivity of the xylem (Ks) and maximum stomatal conductance were both related to the resistance-acquisition gradient in our study system (Bongers *et al.*, 2021; Kröber *et al.*, 2014). Due to the orthogonality of the resistance-acquisition and stomatal control trait gradient in our study system (Fig. 1B), we interpret stomatal control as the extent to which early or late stomatal closure protects the tree's xylem from cavitation during drought at constant levels of cavitation resistance. We thus view stomatal control as a gradient capturing the trade-off between water-spending (i.e. continued transpiration under drought) and water-saving stomatal control (i.e. stomatal closure as protection against cavitation). This view aligns with contemporary perspectives (Martínez-Vilalta & Garcia-Forner, 2017) and fits the species responses we observed in wet to dry years.

However, some association between resistance-acquisition and stomatal control traits may be expected in general, as stomata regulate leaf water potentials to avoid xylem cavitation (McDowell *et al.*, 2008). For instance, there is evidence that $\Psi_{50}$ and the LES are associated with turgor loss point (arguably a proxy for stomatal control) and with (an-)isohydry across species (Fu & Meinzer, 2019; Klein, 2014; Zhu *et al.*, 2018). In contrast, recent local studies did not find any relationship between turgor loss point and $\Psi_{50}$ (Laughlin *et al.*, 2020), nor between turgor loss point and LES traits (Maréchaux *et al.*, 2020). Therefore, the relationships between drought-tolerance traits likely depend on the geographical extent of the study and the range of traits considered. Particularly the multitude of approaches to quantify stomatal control and recent criticisms of classical approaches (Martínez-Vilalta & Garcia-Forner, 2017) may limit our ability to draw general conclusions on the nature and interrelation of both gradients. For instance, stomatal control defined as leaf water potential regulation (i.e. the classical (an-)isohydry definition) has been shown to be not strongly related to leaf gas exchange dynamics or the hydraulic or carbon limitations under drought (Martínez-Vilalta & Garcia-Forner, 2017). In this context, we consider direct measurements of stomatal conductance regulation (or similar ones like sap flux regulation; Schnabel *et al.* (2022)) under gradients of soil and atmospheric drought as crucial to better characterize water-use and drought-tolerance strategies. Universal trait syndromes governing forest responses to drought thus remain controversial (e.g. Guillemot *et al.*, 2022; Henry *et al.*, 2019; Oliveira *et al.*, 2021) and remain a research frontier for future studies.
